# Resolution of *in situ* inflammation in human lupus nephritis into principal immune cell trajectories

**DOI:** 10.1101/2025.02.17.638655

**Authors:** Gabriel Casella, Madeleine Torcasso, Junting Ai, Thao Cao, Satoshi Hara, Michael Andrade, Deepjyoti Ghosh, Anthony Chang, Kichul Ko, Anita S. Chong, Maryellen Giger, Marcus R. Clark

## Abstract

In human lupus nephritis (LuN), tubulointerstitial inflammation (TII) is prognostically more important than glomerular inflammation. However, a comprehensive understanding of both TII complexity and heterogeneity is lacking. Herein, we used high-dimensional confocal microscopy and specialized computer vision techniques to quantify immune cell populations and localize these within normal and diseased renal cortex structures. With these tools, we compared LuN to renal allograft rejection (RAR) and normal kidney. In both LuN and RAR, the 33 characterized immune cell populations formed discrete subgroups whose constituents co-varied in prevalence across biopsies. In both diseases, these co-variant immune cell subgroups organized into the same unique niches. Therefore, inflammation could be resolved into trajectories representing the relative prevalence and density of cardinal immune cell members of each co-variant subgroup. Indeed, in any one biopsy, the inflammatory state could be characterized by quantifying constituent immune cell trajectories. Remarkably, LuN heterogeneity could be captured by quantifying a few myeloid immune cell trajectories while RAR was more complex with additional T cell trajectories. Our studies identify rules governing renal inflammation and thus provide an approach for resolving LuN into discrete mechanistic categories.

Graphical Abstract

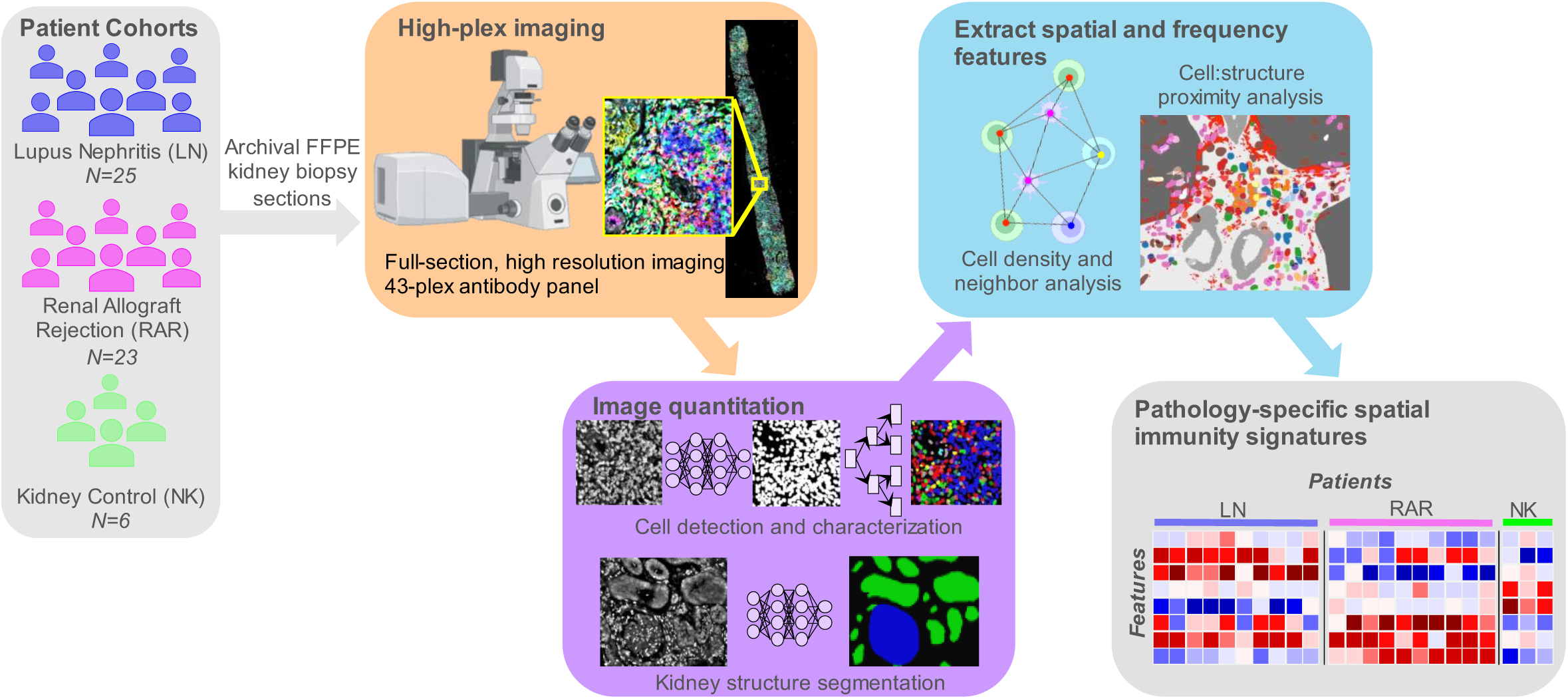

## INTRODUCTION

Among the severe manifestations of systemic lupus erythematosus (SLE), lupus nephritis (LuN) is the most common (*1–5*). Up to 50% of SLE patients develop lupus nephritis, in many cases necessitating treatment with toxic immunosuppressive therapies such as cyclophosphamide or mycophenolate mofetil (*6–8*). Despite such aggressive treatments, many patients do not respond to therapy, and up to 40% of LuN patients progress to renal failure within 5 years of diagnosis (*9–12*).

Much work in human LuN has focused on peripheral autoimmunity and its manifestation within the kidney, glomerulonephritis (GN) (*13*). However, prognosis is more tightly linked to tubulointerstitial inflammation (TII) and scarring (*11, 14–21*). TII, and to some degree GN, are associated with complex *in situ* immune states. Single cell (sc) RNA-Seq from human lupus biopsies has revealed at least 21 different adaptive and innate immune cell clusters (*22*) which organize into complex tubulointerstitial structures ranging from small neighborhoods of CD8^+^ T cells through T:B cell aggregates to canonical germinal centers (*23–27*). TII is associated with selection of B cells expressing unique antibody repertoires and cognate T cell:antigen presenting cell (APC) networks (*24, 28, 29*). These data indicate that LuN is associated with multiple *in situ* immune mechanisms hypothesized to drive local inflammation and tissue destruction (*30*).

Limited studies have related specific *in situ* immune cell populations to prognosis. For example, infiltrating CD8 T cells have been associated with a poor prognosis (*31, 32*). In a study of principal adaptive immune cell populations, densities of CD4-T cells were most predictive of resistance to conventional therapy and progression to renal failure (*27*). Furthermore, intrarenal T cells promote tissue-injury in murine LuN and share effector pathways with kidney-infiltrating T cells in human LuN (*33*). Finally, the efficacy of the calcineurin-inhibitor voclosporin highlights the importance of T cells in some patients with LuN (*34*).

High densities of myeloid cells were associated with progressive disease (*35–37*). However, these studies used simple markers of myeloid cells while recent studies have revealed great heterogeneity in the intrarenal myeloid cell compartment (*22, 38*). It is unclear which *in situ* myeloid population is most closely linked to prognosis in LuN.

The striking responses of lupus patients to CD19 CAR T cells have renewed interest in the pathogenic role of B cells (*39, 40*). So far, clinical trials with CD19 CAR T cells have been small, uncontrolled, and with limited mechanistic studies. Furthermore, they appear at odds with observations that in LuN, *in situ* CD20+ B cell densities are associated with a good prognosis (*27*). These contradictory results could simply reflect different patient populations or the effects of conditioning chemotherapy. However, in aggregate, these observations might suggest unexplored relationships between systemic and *in situ* autoimmunity and/or *in situ* functions(s) for B cells not captured by simple total cell densities (*41, 42*). These functions might include secretion of highly pathogenic antibodies or sub-populations of B cells presenting antigen to large populations of pathogenic T cells (*43, 44*).

The above studies have each focused on the potential pathogenic role of individual cell populations. There has been no comprehensive study of the relationships between these immune cell populations and how their frequencies and spatial distributions differ between LuN patients. The latter is likely important as how adaptive immune cells organize in the LuN kidney provides unique prognostic information (*27*). Herein, we used high-dimensional confocal microscopy and customized computer vision tools to capture *in situ* LuN immune cell frequency and spatial distribution heterogeneity and compare these to those of renal allograft rejection (RAR). Our data suggest that *in situ* LuN and RAR heterogeneity can be described by quantifying a limited number of shared and unique immune cell states. Furthermore, our data suggest that each immune state develops along partially mutually exclusive trajectories the constituents of which are associated with specific features of renal inflammation and damage.

## RESULTS

### Lupus nephritis and renal allograft rejection patient biopsies

To probe the *in situ* immune differences between prototypical autoimmune (LuN) and alloimmune (RAR) renal diseases, we acquired 25 LuN and 23 RAR initial diagnostic biopsies. An additional six normal kidney control (KC) samples were acquired from nephrectomies for renal cell carcinoma. In LuN, tubulointerstitial inflammation (TII) is more prognostically important than GN (*11, 14–19*) while RAR is primarily a tubulointerstitial disease (*45*). Therefore, our analysis focused on comparing TII between both diseases. Conventional histological scoring by a pathologist of TII, chronicity, interstitial fibrosis, and tubular atrophy did not reveal any significant differences between the two disease cohorts (**Fig. 1A**). **Supplemental Figure 1A** provides LuN class, patient age at diagnosis and age at biopsy. **Supplemental Figure 1B** provides information on RAR patients including rejection subtype, donor type, age at transplant and age at time of biopsy.

**Figure 1.**
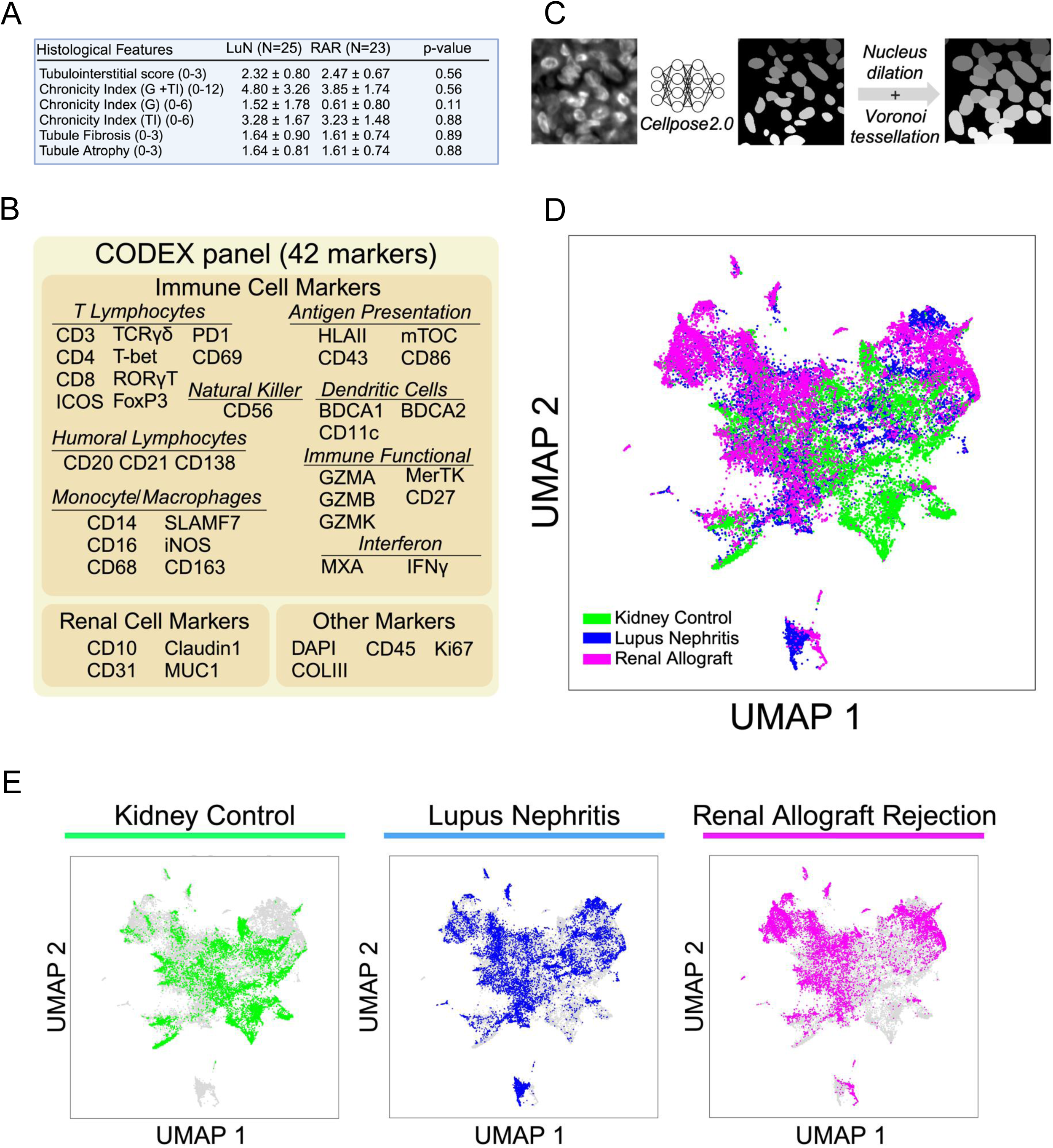
Cell detection in multiplex microcopy imaging of renal biopsy tissue. A. Histological scoring of patient paired H&E and PAS biopsies. The mean autoimmune cohort and standard deviation of the histological features are shown. Mann-Whitney-U nonparametric difference in means p-values are shown. B. CODEX antibody marker panel grouped by cell lineage or cellular activity most associated with that marker. C. Workflow diagram of cell detection and segmentation. Cell nuclei are defined using Cellpose 2.0 on a 512×512 DAPI image after performing Human-in-the-loop model retraining. Cell body was subsequently approximated by performing nuclear dilation with Voronoi tessellation. 512×512 images of DAPI segmentations are then knitted and aligned with the 42 fluorescent channels. D. UMAP dimensional reduction of cell body MFI from 30,000 cells randomly sampled. 10,000 cells are sampled from each of the three cohorts: Normal Kidney Control, Lupus Nephritis, and Renal Allograft Rejection. Cells are colored according to their cohort of origin: KC, LuN, and RAR. E. UMAP dimensional reduction of cell body MFI from 30,000 cells randomly sampled. Cells are colored according to their cohort of origin.

### Quantifying highly multiplex images of full biopsy renal sections

We hypothesized that there would be significant differences in *in situ* immune cell populations between LuN and RAR. Therefore, we stained each biopsy with antibodies specific for 42 markers that identified known *in situ* immune cell constituents in LuN and RAR (*22, 46*), other important cell populations (e.g. ψ8 T cells), proximal tubules (CD10), distal tubules (mucin 1, MUC1) as well as markers of inflammation (Myxovirus resistance protein 1, MXA) and scarring (collagen III, COLIII)(**Fig. 1B**).

For our imaging, we used spinning disk confocal microscopy coupled to a co-detection by indexing (CODEX) microfluidic head and CODEX chemistry (*47*). We captured whole slide images with a pixel size of 0.1507μm. Each cycle of imaging included a nucleus stain (DAPI) imaged at 405nm and four other stains imaged at 488nm, 561nm, 637nm and 730 nm. In our analytic pipeline, we adapted ASHLAR (*48*) to stitch and align image tiles into full-sections to generate an accurate coordinate space. Background was subtracted and the resulting full biopsy image stacks were min-max normalized to the 99th percentile for each channel.

Analysis of the resulting large and complex datasets required first detection of cells and approximation of cell boundaries (instance segmentation) and then assignment to known cell class (annotation). For nuclear segmentation of DAPI-stained renal tissue, we initially tested several convolutional neural networks (CNNs) including Mask R-CNN (*49*), Cellpose 2.0 (*50*), Deepcell (*51*) and Stardist (*52*). However, qualitative comparisons to manually annotated images revealed Cellpose to perform best, especially in highly inflamed areas (data not shown). Therefore, Cellpose 2.0 was incorporated into our analytic pipeline (**Fig. 1C**).

To assess cell detection and segmentation performance, we compared the Cellpose output to a manually segmented dataset of 100 regions of interest (ROIs) from five LuN biopsies and five RAR biopsies (ten ROIs/biopsy). Cellpose had a zero-shot F1 performance score of ∼0.80 (LuN) and ∼0.58 (RAR), with an average precision of ∼0.67 and ∼0.38 respectively at an intersection over union (IoU) of 0.25 (**Supplemental Fig. 1C-D**). When we adopted human-in-the-loop (HITL)(*53*) fine-tuning, we achieved a F1 performance score of ∼0.88 (LuN) and ∼0.73 (RAR) with an average precision of ∼0.77 and 0.53, respectively. Approximately 2.19 million cells were detected across all 54 samples.

The Cellpose DAPI nuclear segmentation mask was dilated approximately 1 micron with Voronoi tessellation to capture cytoplasmic staining and approximate whole cell body boundaries. From this cell body mask, we captured mean fluorescence intensity (MFI) which was standardized across all 42 channels. We used uniform manifold approximation and projection (UMAP) for dimension reduction and for plotting a random 10,000 cells from each patient cohort in two-dimensional space (**Fig. 1D and E**).

The distribution of cells for each patient cohort was different. All three cohorts had similarities and difference in their distributions within the UMAP space. To begin to understand these differences, we projected the distributions of all 42 markers onto the UMAP space (**Supplemental Fig. 2A-B**). Several lymphocyte markers, including CD3, CD4, CD8 and CD20 strongly co-localize in the upper left of the UMAP plots. In contrast, some myeloid immune cell markers, such as CD14 and BDCA1, co-localize in both the upper left and right quadrants. These data suggest that infiltrating lymphocytes reside in the upper left quadrant while myeloid populations localize in the upper left and right quadrants.

It was apparent from Supplemental Fig. 2 that FOXP3 and other markers had apparent broad distributions. Furthermore, in areas of dense inflammation, fluorescence signals can bleed into proximate cells.

Finally, because we are randomly sampling a two-dimensional cut of a three-dimensional object, the staining intensity within a given cell class can be variable. For these reasons, identifying different cell populations by K-means clustering in the UMAP space, as is done for scRNA-Seq data, is not adequate (*54*).

To circumvent these limitations, we emulated the hierarchical approach used to immunophenotype cells by flow-cytometry. Briefly, the mean pixel intensities for 26 cardinal markers were organized into decision trees (DTs) for cell class annotation (*47, 55*)(**Supplemental Fig. 3**). In **Supplemental Fig. 3A** is given the DT for CD45+ immune cells. Some macrophage populations expressed undetectable levels of CD45 (Supplemental Fig. 2A). Therefore, CD45-renal structures and macrophage populations were assigned as in **Supplemental Fig. 3B and 3C**. T cell subtypes were assigned as in **Supplemental Fig. 3D**. The resulting 33 annotated cell populations are shown in **Supplemental Fig. 4A**. In this way, markers such as FOXP3 were not used for global assignment decisions. Rather, FOXP3 was used to identify regulatory cells within T cell populations.

Approximately 77% of segmented cells were assigned to a cell class across disease groups. Primarily renal tubular cells could not be assigned to a class as identifying markers often stained the plasma membrane which was beyond the one-micron DAPI dilation in these large cells. This limitation was circumvented as described below.

Each annotated cell class expressed the expected cardinal markers (**Supplemental Fig. 4B**). Furthermore, across biopsy cohorts, each annotated class manifested similar distributions of cell staining (**Supplemental Fig. 4C**). These data suggest quantitative comparisons could be made across patient cohorts.

Using the above DTs to annotate cells in the UMAP space, similarities and differences between biopsy cohorts became apparent (**Fig. 2A**). While renal parenchymal structures occupied the center of all three cohort UMAPs, there were differences in distributions. Some of these changes could be ascribed differential expression of markers of inflammation (MXA, Claudin 1) and scarring (COLIII) in the disease cohorts (**Supplemental Fig 4D**). Lymphocytes, especially T cells, occupied the upper left quadrant. These populations were scant in normal kidney and highest in RAR. In contrast, myeloid populations were increased in both LuN and RAR. However, there were some differences in the distribution of myeloid cells in the UMAP space suggesting that each disease might be enriched for specific populations.

**Figure 2.**
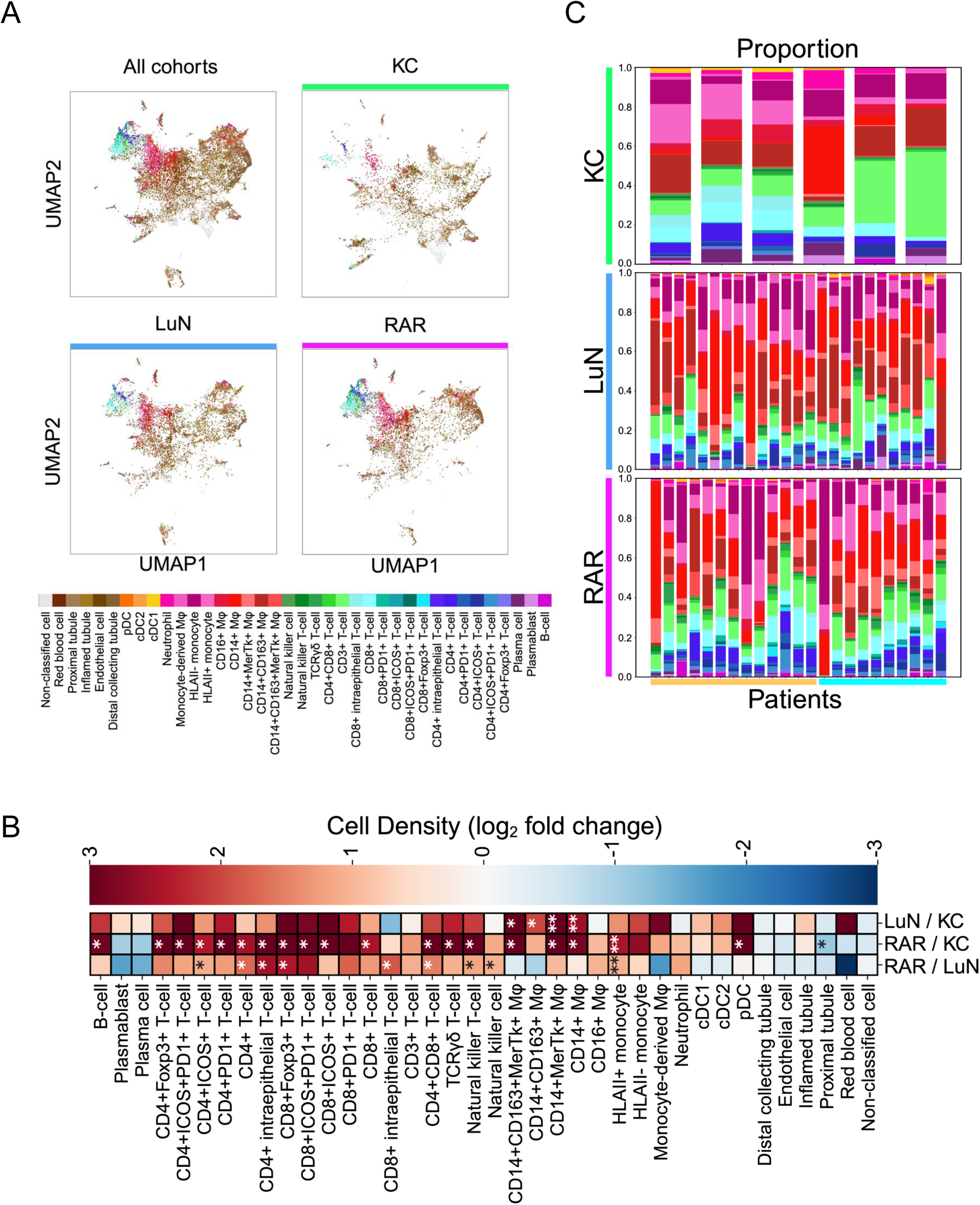
Distribution of immune cell classes in KC, LuN and RAR. A. UMAP dimensional reduction of cell body MFI from 30,000 cells randomly sampled. 10,000 cells were sampled from each of the indicated cohorts. B. Non-parametric Mann-Whitney-U difference of the mean test for population differences in classified cell density between patient cohorts; from the top row: LuN-KC, RAR-KC, and RAR-LuN. Color indicates log2 fold change. Benjamini-Hochberg p-value correction was performed. * p < 0.05, ** p < 0.01, *** p < 0.001. C. Patient-level proportions of the 33 immune cell classes by cohort. Kidney control (*top*), lupus nephritis (*middle*), and renal allograft rejection (*bottom*). Mixed-rejection biopsies are denoted using a beige bar, T-cell mediated rejection biopsies are denoted in light blue. All cell classes (except non-classified cells) are color-coded as displayed in the panel in Figure 2A. Bars at bottom identify MR (yellow) and TCMR (blue).

A more detailed picture was provided when constituent cell percentages were examined (**Supplemental Fig. 5A**). As expected, renal parenchymal cells were the most prevalent. As described above, these were under estimations. Among immune cell populations, myeloid cells, especially CD14+ macrophages and inflammatory monocytes, were enriched in both LuN and RAR. However, total CD163+ macrophages were enriched in LuN. In contrast, most T cell populations, including CD4+, CD8+ and ψ8 T cells, were enriched in RAR. Indeed, comparing cell densities between KC, LuN and RAR revealed that LuN was enriched for myeloid cell populations, especially CD163+ macrophages (**Fig. 2B**). In contrast, RAR was enriched for multiple T cell populations. Notably CD3+ T cells lacking CD4, CD8 or γ co-expression (double negative or DN T cells), were not enriched in either disease compared to KC (*27, 56*). These data suggest that, in our cohort, RAR is characterized by enrichment of T cell and myeloid populations while LuN is enriched primarily in myeloid cell populations.

In comparison to KC biopsies, there was substantial heterogeneity in disease groups both in terms of lymphocyte versus myeloid cell populations in each biopsy and the constituents within each broad immune cell class (**Fig. 2C**). In LuN, there were four to five biopsies with relatively high lymphocyte populations while in the rest myeloid cells predominated. Myeloid cells predominated in about half of RAR biopsies. Furthermore, especially in RAR, the myeloid compartment was dominated by either macrophages, or HLA class II positive or negative inflammatory monocytes (defined by CD14 and CD16 expression)(*57*). Among the RAR samples were 13 mixed rejection (MR) and ten T cell mediated rejection (TCMR). However, there were no apparent, global differences in the distributions of lymphoid or myeloid cells between these disease classes. As these subgroups were small, no further comparisons between the two were performed.

### Distinct immune trajectories in LuN and RAR

Plotting Spearman’s correlation between immune cell proportions across LuN and RAR biopsies revealed co-variance between specific subpopulations. (**Fig. 3A, B, Supplemental Fig. 6A, B**). In both diseases, co-variance across T cell populations, including CD4+, CD8+ and ψ8 T cells, was the most striking. Interestingly, there was not a strong correlation between B cells and either plasma cells or plasmablasts and in neither disease was there a correlation between plasma cells/blasts and most T cell populations. Focusing on innate cells, there were co-variant subpopulations or blocks among the most common cell populations in both LuN and RAR. Specifically, we observed co-variance between macrophages expressing CD163 including CD14+CD163+MerTk+ and CD14+CD163+ populations. There were separate co-variant cell blocks for CD163-CD14+ macrophages (+/- MerTK) and for CD14+CD16+ monocytes (+/- HLA class II).

**Figure 3.**
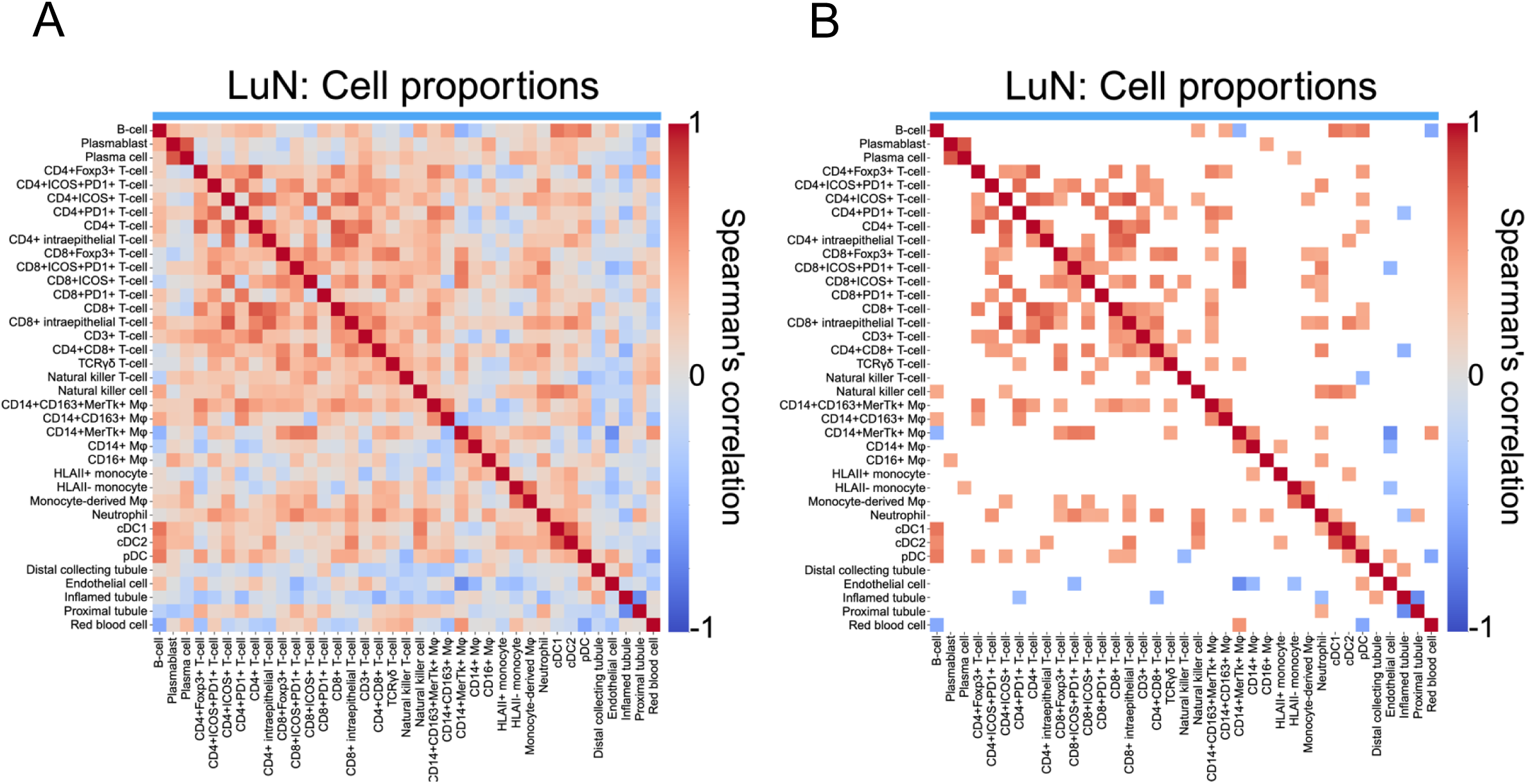
Spearman’s correlation of lupus nephritis cell class proportions. A. Heatmap of the non-parametric spearman’s correlations between patient-level immune and non-immune cell proportions for lupus nephritis patients. B. Heatmap of statistically significant (p<0.05) non-parametric spearman’s correlations between patient-level immune and non-immune cell proportions for lupus nephritis patients.

In RAR, some of the above populations had inverse relationships. For example, CD163+ macrophages inversely varied with some CD163-macrophage and monocyte populations. In LuN, there was a trend towards these inverse relationships. In LuN, B cell densities have been associated with a good prognosis (*27*). Therefore, it is interesting that in LuN, they had an inverse proportional relationship with CD14+MerTK+ macrophages and a positive correlation with some CD163+ macrophage populations, both of which are much more prevalent than B cells. Similar correlations were observed when considering the densities of the different cell populations (**Supplemental Fig. 6C-F**) suggesting that the observed relationships were robust. These separate, co-variant subpopulations, or cell blocks, suggest that a limited number of discrete immune states characterize both LuN and RAR.

Cohort specific relationships were observed when we plotted the densities of all myeloid cells versus all T cells for each biopsy (**Fig. 4A**). The size of each point reflects total humoral cells. Strikingly, LuN unfolded along a myeloid axis and RAR along a T cell axis with some RAR biopsies also having substantial densities of myeloid cells. These differences in myeloid and T cell densities were associated with striking visual differences (**Fig. 4B).** We then plotted the densities of CD14+MerTk+CD163-macrophages versus CD14+CD163+ macrophages versus total CD8+ T cells (**Fig. 4C**). Interestingly, RAR unfolds along the CD14+MerTk+CD163-axis which is also rich in CD8+ T cells. In contrast, some LuN biopsies also had relatively high CD163-macrophage densities (4/25 biopsies) while other biopsies had high densities of CD163+ macrophages (5/25).

**Figure 4.**
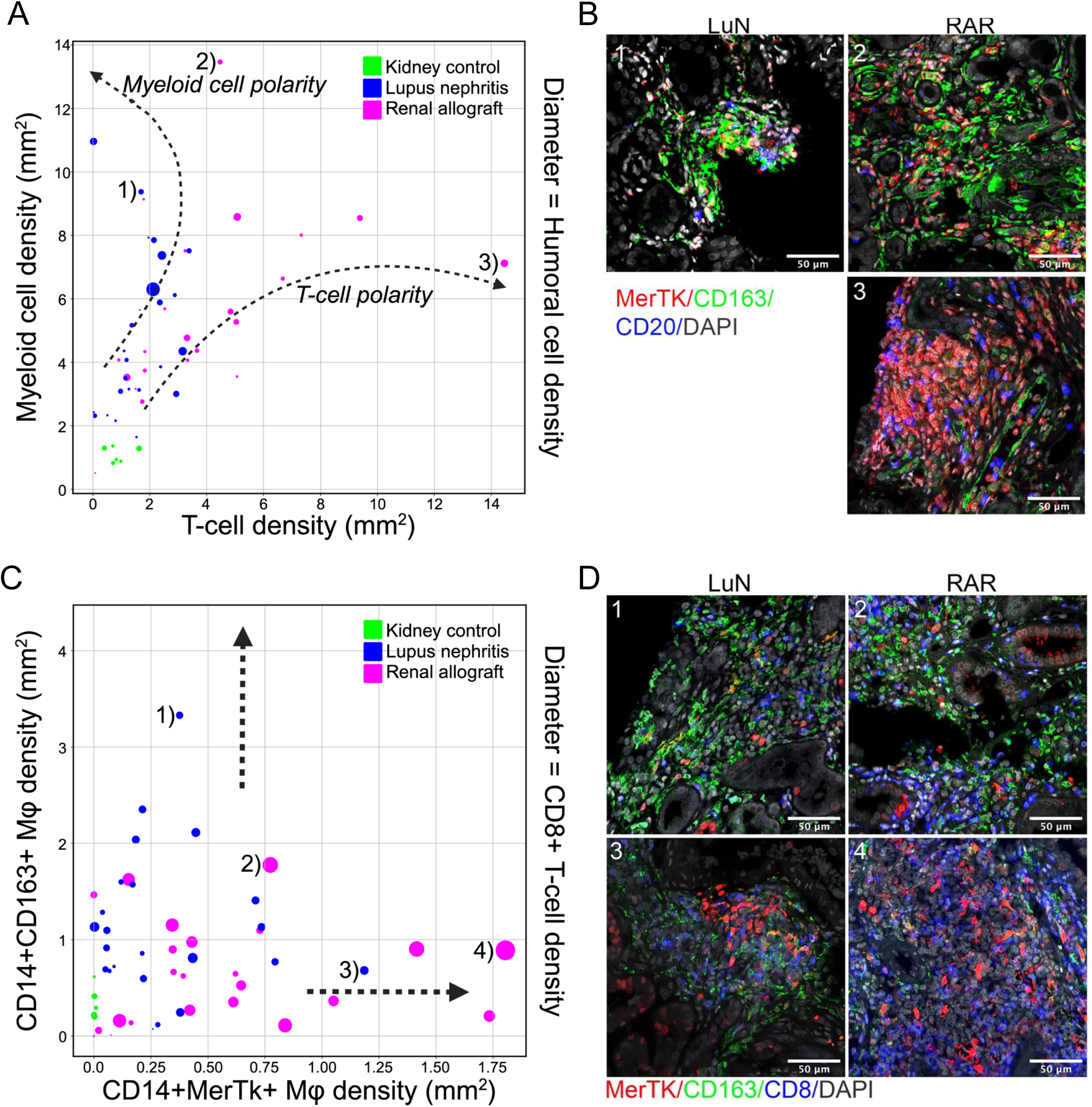
Distinct immune trajectories are associated with distinct pathologic states. A. Plot of the patient-level T-cell density (*x-axis*) and myeloid cell density (*y-axis*) colored by cohort: Kidney control (*green*), LuN (*blue*), RAR (*magenta*). Diameter indicates humoral cell density. B. Representative microscopy images of immune cell polarization in lupus nephritis and renal allograft rejection patient cohorts. Image numbers correspond to biopsies indicated in “A”. C. Plot of the patient-level CD14+MerTk+ macrophage density (*x-axis*) and CD14+CD163+ macrophage density (*y-axis*) colored by cohort: Kidney control (*green*), LuN (*blue*), RAR (*magenta*). Diameter indicates T-cell density. D. Representative microscopy images of CD14+CD163+ and CD14+MerTk+ enriched biopsies. Image numbers correspond to biopsies indicated in “C”.

Examples from the indicated biopsies are provided in **Fig. 4D**. These data suggest that in our patient cohorts, LuN and RAR biopsies often lie along different immune cell trajectories.

### Immune cell localization within renal compartments

The above analyses examined immune cell frequencies across whole biopsies. However, the kidney is structurally complex with glomeruli, tubules and the tubulointerstitial space. Compared to hematopoietic cells, tubules have a characteristic pattern of DAPI staining. Therefore, to identify tubules, Omnipose (*58*) was trained on representative 10-fold downsized DAPI kidney images from CODEX image stacks. Glomeruli were segmented manually (**Supplemental Figs. 7A-B**).

We first assessed the densities of all immune cell populations across the different renal compartments (**Fig 5A**). From left to right are provided interstitial, tubular, glomerular, peritubular (dilated tubular mask) and periglomerular (dilated glomerular mask) densities for all indicated immune cell populations. In both diseases, and for all immune cell populations, densities were highest in the periglomerular space. In general, there were more plasma cells/blasts in LuN distributed across all compartments. In contrast, RAR was generally enriched in T cells distributed across all renal structures except for glomeruli. Furthermore, there was a general enrichment of CD163+ macrophages in LuN.

**Figure 5.**
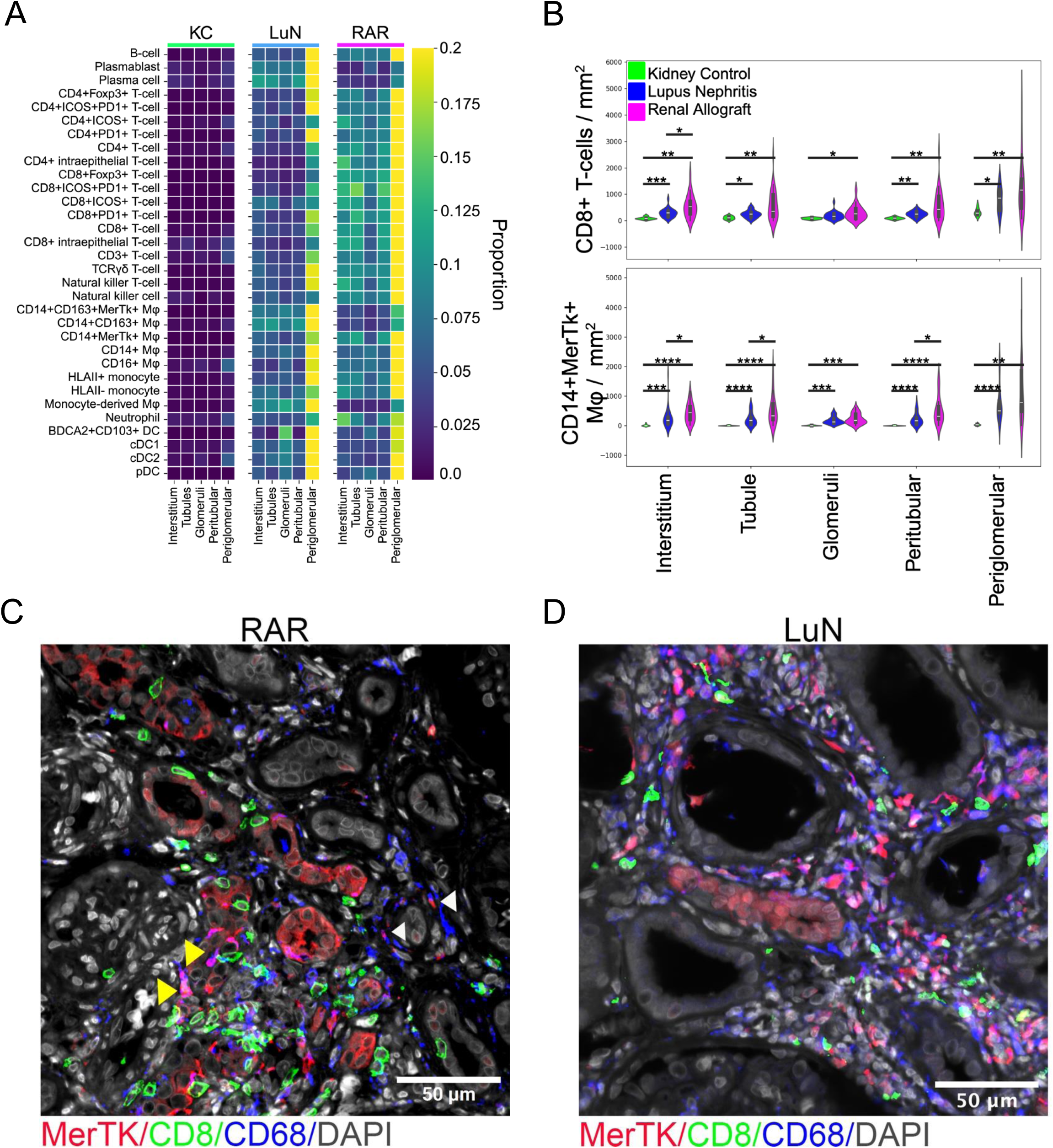
Distribution of inflammation across renal compartments. A. Heatmap showing the biopsy-average density of immune cells of the kidney anatomical compartments, normalized to proportion across cohorts. From the left: interstitium, tubule, glomeruli, perititubular, periglomerular. (*Left heatmap*) kidney control, (*middle heatmap*) lupus nephritis, and (*right heatmap*) renal allograft respectively. B. Violin plots of the biopsy-average number of cells per mm^2^ within the indicated tissue compartments for CD8+ T-cells and CD14+MerTk+ macrophages (Mϕ). Mann-Whitney U test. *<.05, **<.01 and ***<.001. C. RAR example demonstrating tubulitis (yellow arrowhead) and peri-tubular inflammation (white arrowhead). D. LuN example demonstrating diffuse interstitial inflammation.

We then focused on two potential disease-associated populations, CD8+ T cells and CD14+MerTK+ macrophages (**Fig. 5B**). In both cell populations, the peri-glomerular enrichment was evident. In addition, in some RAR biopsies there is an enrichment of both cell populations in the peritubular and tubular space compared to the interstitial space. Indeed, in RAR biopsies there was both peritubular inflammation and tubulitis (**Fig. 5C**). In contrast, interstitial and peritubular densities were similar in LuN and tubulitis was rare (**Fig. 5D** and data not shown). These data suggest that inflammation in the RAR tubulointerstitium is centered around and in tubules while in LuN it occurs diffusely through the interstitium.

### Organization of inflammation into neighborhoods

We next used DBSCAN (*59*) to determine if *in situ* inflammation was organized in LuN and RAR. K-means clustering and bootstrapping were first used to estimate the optimal number of states using spatial coordinates for all immune cells and the indicated cell neighborhood size exclusion conditions (**Sup. Fig. 8**). We found that the best fit K-clusters were consistently between 10 and 11. We used 11 as the optimal number of neighborhoods for downstream analysis.

We generated a heatmap using features including average total cell count across biopsies, and average cell proportion to visualize the unique elements of each neighborhood (**Fig. 6A**). When cell counts were examined, neighborhoods 1, 3, 4, and 7 had the most cells, while neighborhoods 5, 0, 8 and 9 were the most frequent (**Fig. 6B**). The most frequent cell neighborhoods had unique features. Notably, was the enrichment of CD14+CD163+ macrophages and CD14+MerTk+ macrophages in neighborhoods 0 and 5 respectively.

**Figure. 6.**
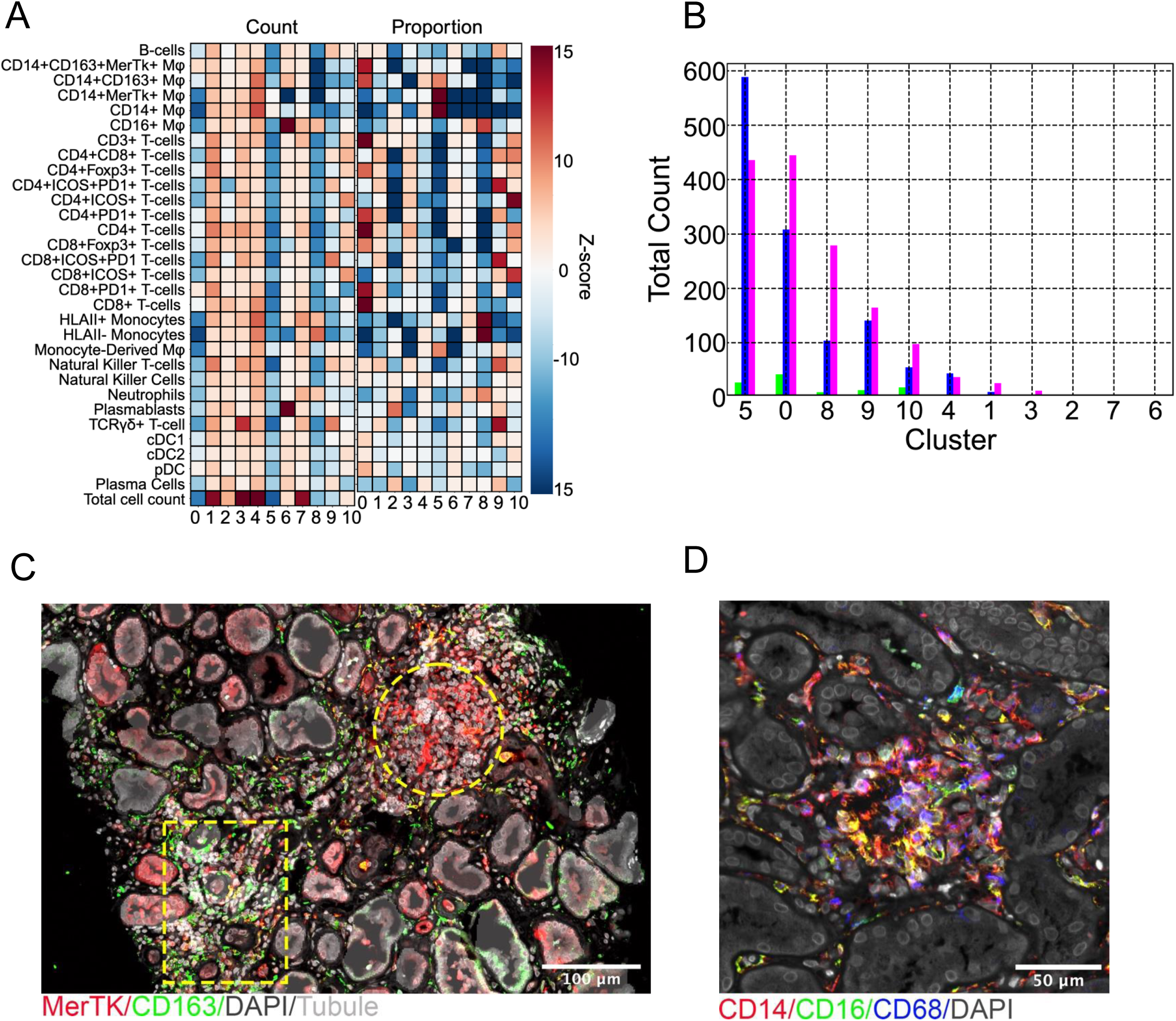
Distribution of cell classes in in situ immune clusters. A. Heatmap of the Z-score of Extracted Features from DBSCAN cell networks using an optimal K=11 means clustering of cell clusters larger than 19 cells. (Left) Cell count of the immune 30 cell types classified and the total cell count in the cell networks, (right) proportion of the same immune cells in the cell networks. B. Bar plot showing the total count of the various K=11 DBSCAN cell networks colored by patient cohort:: KC (green), LuN (blue), and RAR (magenta). C. Example of clusters 0 (box) and 5 (circle) within the same biopsy. D. Example of cluster 8.

Neighborhood 8 was enriched in inflammatory monocytes and CD16+ macrophages while neighborhood 9 contained noncanonical T cell populations including ψ8 T cells and NK T cells. Representative examples of neighborhoods 0 and 5 are shown in one biopsy indicating that clusters of CD163+ and CD163-macrophages can occur in the same biopsy (**Fig. 6C**). An example of the inflammatory monocyte enriched neighborhood 8 is provided in **Fig. 6D**.

### Immune cell correlates with tissue inflammation and damage

From our high-dimensional staining panel, we could derive four measures of tubulointerstitial inflammation and scarring: 1) total biopsy immune cell density. 2) total biopsy inflamed tubule cell density (MXA and Claudin 1 in tubule mask). 3) COLIII mask area (% of total biopsy area). 4) MXA mask tissue area (% of total biopsy area). Given the above findings, we plotted these four measures versus whole biopsy densities and proportions of total CD8+ T cells, total CD4+ T cells, CD14+MerTK+CD163-macrophages, CD14+CD163+ macrophages, HLA class II+ inflammatory monocytes and HLA class II-inflammatory monocytes, and then performed ordinary least squares linear regression (OLS). Only graphs with positive correlations are provided.

We observed that CD8+ T cell densities were correlated with immune cell densities in RAR and LuN (**Fig. 7A**). In contrast, only in RAR were CD8+ T cell densities associated with MXA expression (**Fig. 7B**). Consistent with the co-variance of CD4+ and CD8+ T cell densities across biopsies, similar associations were observed for CD4+ T cell densities (**Figs. 7C-D**). CD14+MerTk+CD163-macrophage cell densities were associated with immune densities in both diseases (**Fig. 7E**). CD14+MerTk+ macrophages were also associated with inflamed tubule density in RAR with a similar trend in LuN (**Fig. 7F**). There was a strong association between CD163+ macrophage cell densities and immune cell densities in LuN but not RAR (**Fig. 7G**).

**Figure 7:**
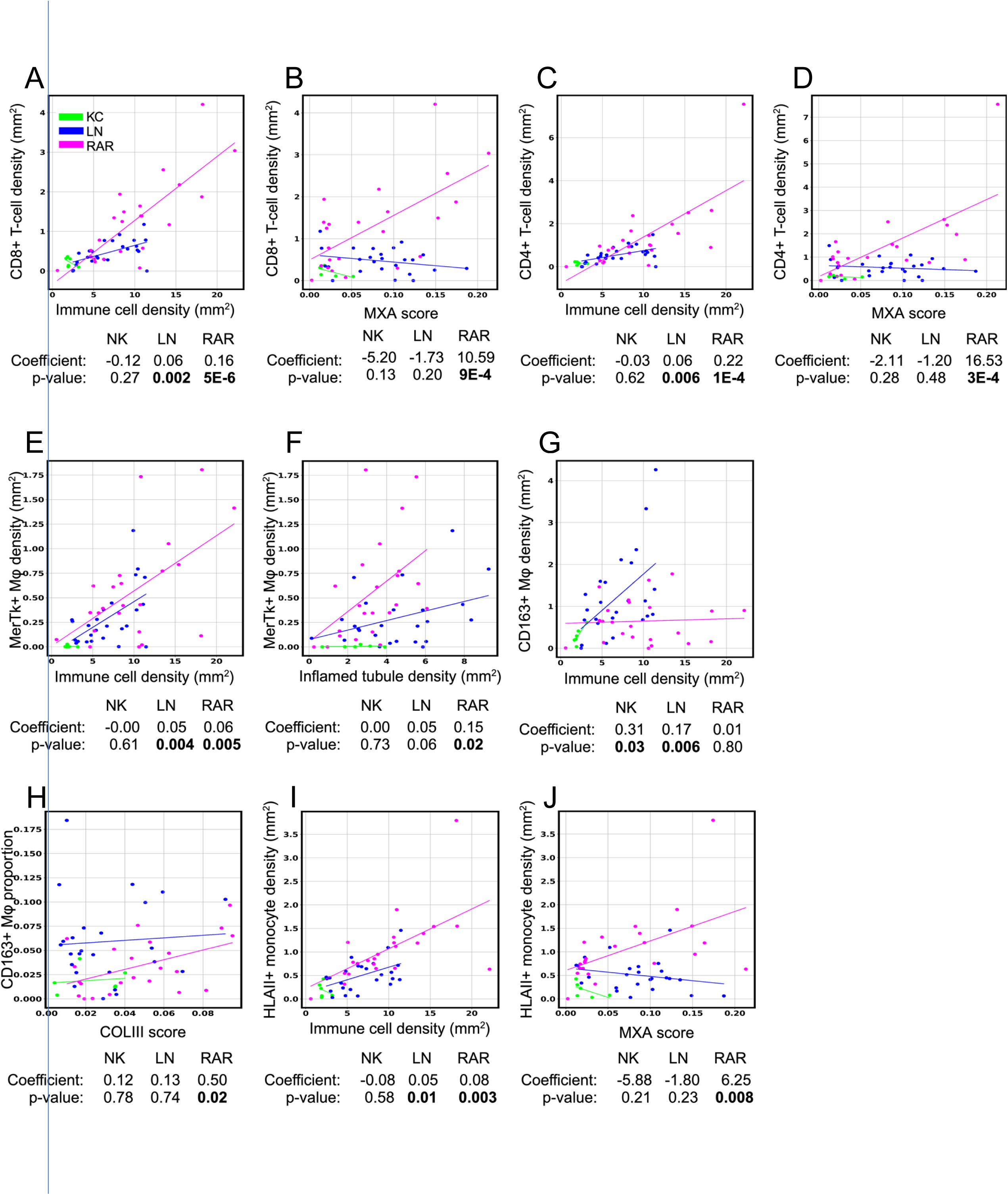
Immune cell trajectories correlate with tissue inflammation and damage. A-B. CD8+ T-cell density as a function of immune cell density (A) and score (B). C-D. CD4+ T cell density as a function of immune cell density (C) and myxovirus resistance protein 1 (MXA) score (D). E-F: MerTK+CD163-macrophage density as a function of immune cell density (E) and inflamed tubule density (F). A. CD163+ macrophage density as a function of immune cell density. B. CD163+ macrophage proportion as a function of Collagen III (COL III) score. JI-J. Inflammatory HLA class II+ monocytes immune cell density as a function of immune cell density (I) and MXA score (J).

However, CD163+ macrophage proportions were associated with COL III scores only in RAR (**Fig. 7H**). Finally, densities of HLA class II+ inflammatory monocytes were associated with immune cell densities (**Fig. 7I**) in both diseases. However, they were only associated with MXA scores in RAR (**Fig. 7J**). These data indicate that in LuN and RAR, immune cell populations having the same surface phenotype can have either similar or different associations with measures of renal inflammation and scaring.

### Immune cell trajectories characterize individual biopsies

The above data suggest that inflammation heterogeneity can be resolved into relatively few co-variant blocks of the most prevalent immune cells. Immune cell constituents of these different inflammatory states organized into distinct niches and were associated with specific manifestations of renal inflammation and scarring. To begin to graphically quantify inflammation in a way that could be compared across diseases and biopsies, we generated radar diagrams in which each axis was the density of a principal cell population: total CD8+ T cells, total CD4+ T cells, CD14+CD163-macrophages, CD163+ macrophages, HLA class II+ inflammatory monocytes, and HLA class II-inflammatory monocytes.

We first plotted all 54 individual biopsies on a single radar graph, color coded by clinical cohort (**Fig. 8A**). Compared to both LuN and RAR, the KC samples had far less densities of these six immune cell population groups. It is also apparent that some RAR biopsies had higher densities of several immune cell populations compared to LuN biopsies. Indeed, plotting average immune cell densities for each clinical cohort indicate that all populations, except for CD163+ macrophages, are higher in RAR than LuN (**Fig. 8B**). These data indicate that, on average, the RAR biopsies are more inflamed even though the two disease cohorts were scored similarly for TII (tubulointerstitial score) by a renal pathologist (Fig. 1A).

**Figure. 8.**
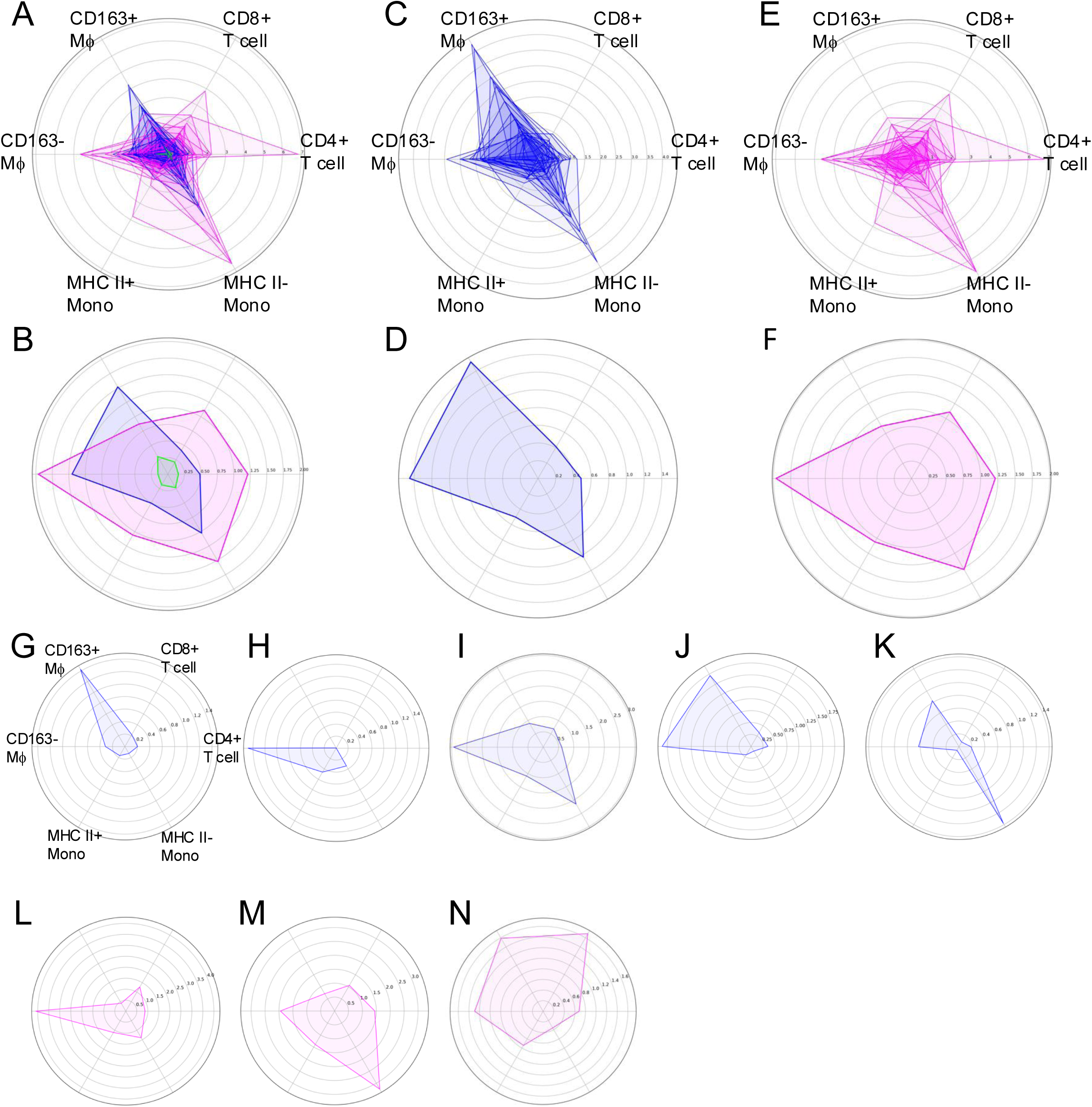
Quantifying *in situ* immune state using principal immune cell trajectories. A. Composite web graph of all individual biopsies colored by cohort. Green: normal kidney, blue: LuN and purple: RAR. B. Plot of averages for each clinical cohort. C. Plot of LuN individual biopsies. D. Average LuN densities for indicated immune cell populations. E. Plot of RAR individual biopsies. F. Average RAR densities for indicated immune cell populations. G. Example of individual LuN biopsy with CD163+ macrophage polarity. H. Example of LuN biopsy with CD163-macrophage polarity. I. Example of LuN biopsy with CD163-macrophage and HLA II inflammatory monocyte polarities. J. Example of LuN biopsy with CD163+ and CD163-macrophage polarity. K. Example of LuN biopsy with CD163+ and CD163-macrophage polarity in combination with HLA II-inflammatory monocyte polarity. L. Example of individual RAR biopsy with CD163-macrophage polarity. M. Example of individual RAR biopsy with multiple immune cell lineages plus HLA II-inflammatory monocytes trajectory. N. Example of individual RAR biopsy with multiple immune cell lineages without HLA II-inflammatory monocytes trajectory.

Plotting LuN and RAR biopsy cohorts separately allowed better visualization of the immune cell densities that characterized each disease cohort. In LuN, inflammation unfolds along three major density axes, CD163+ macrophages, CD163-macrophages and HLA Class II-inflammatory monocytes (**Figs. 8C-D**). Across the disease cohort, RAR appears more complex with significant projections along the CD163-macrophage, HLA Class II+ inflammatory monocytes, HLA Class II-inflammatory monocytes, CD4+ T cells and CD8 T cell density axes (**Figs. 8E-F**). Notable is the lack of substantial CD163+ macrophage densities. These data suggest that LuN is characterized by a few myeloid immune cell density axes while RAR has both myeloid and adaptive immune cell density axes.

We then plotted individual disease biopsies. Of the 25 LuN biopsies, six were characterized by prominent CD163+ macrophage densities (>2-fold over CD163-macrophages) and a relative lack of other immune cell populations (**Fig 8G, Supplemental Fig. 9A**). One of these biopsies also had high numbers of HLA class II-inflammatory monocytes (**Supplemental Fig. 9B**). Similarly, six biopsies were characterized by primarily CD163-macrophage populations (**Fig 8H, Supplemental Fig. 9C**). Two of these also had substantial HLA class II-monocyte immune cell densities (**Fig. 8I, Supplemental Fig. 9D)**. The remaining 13 biopsies had both substantial CD163+ and CD163-immune cell densities. Seven of these biopsies did not have substantial densities of other populations (**Fig. 8J**, **Supplemental Fig. 9E**). The other six had concurrent HLA class II-inflammatory monocyte densities (**Fig. 8K** and **Supplemental Fig. 9F**). Therefore, within our cohort, LuN fell into well-defined subsets characterized by CD163+ macrophage, CD163-macrophage and HLA class II-inflammatory monocyte densities.

RAR was more complex. Of the 23 biopsies, eight had a predominantly a single immune cell population (> two-fold difference over other immune cell populations). In five of these, CD163-macrophage densities predominated (**Fig. 8L**, **Supplemental Fig. 10A**). In two, HLA class II-inflammatory monocytes predominated and in one, CD163+ macrophages. Only two biopsies were characterized by two trajectories, both of which included CD163-macrophages (**Supplemental Fig. 10B**). The rest manifested multiple myeloid and adaptive immune cell densities including six that had HLA class II-inflammatory monocytes (**Fig. 8M, Supplemental Fig. 10C**) and seven that did not (**Fig. 8N, Supplemental Fig. 10D**). These data suggest that, in our cohort, RAR falls broadly into two categories, those that can be characterized by myeloid cells, most commonly CD163-macrophages, and those characterized by both myeloid and adaptive immune cell densities.

## DISCUSSION

Using high dimensional imaging and computer vision techniques specifically adapted for the kidney, we provide the first comprehensive assessment of LuN and RAR immune cell constituency, how these cells are organized into neighborhoods and their relationships to renal cortical structures. These data resolve immune cell heterogeneity into a limited number of cell states. Indeed, in any one biopsy, the inflammatory state could be characterized by the relative prevalence and magnitude of cardinal immune cell trajectories. Notably, in many biopsies from both diseases, CD163-macrophages predominated with or without other concurrent immune cell populations. In LuN, concurrent immune cell populations were of myeloid lineages while in RAR CD163-macrophages co-occurred with both myeloid and adaptive immune cell lineages. While both diseases manifested inflammatory states anchored by CD163-macrophages, only in LuN there were several biopsies in which CD163+ macrophages predominated. Our studies suggest an approach to quantifying *in situ* immunity that allows comparisons between individual biopsies in and across disease states.

Canonical cell markers identified populations of immune cells that co-varied with each other and shared common spatial distributions within the tubulointerstitium. CD163+, a marker of some M2 macrophages, identified cell populations that formed neighborhoods and distributions different from other macrophage populations regardless of other shared markers such as the phagocytic receptor MerTK (*60*). Likewise, inflammatory monocytes, did not co-vary with any other immune cell blocks/subgroups and formed distinct neighborhoods (*57*). Our current staining panel did not allow fine categorization of either macrophage or monocyte subsets.

Within the limitations of these small patient cohorts, both CD4+ and CD8+ T cells largely behaved as a covariant block and co-segregated into specific neighborhoods. Remarkably, these fundamental relationships were apparent in both LuN and RAR. Rather, these cell subgroups and niches primarily varied in relative prevalence between the two diseases. T cell blocks were a dominant feature of RAR while in LuN macrophage subgroups were more prevalent. These stereotypic relationships between functionally related immune cell populations suggest that, in individual patients, inflammation develops along a limited number of trajectories.

While LuN and RAR can manifest the same immune cell trajectories, they were often associated with different features of tubulointerstitial inflammation and damage. For example, only in RAR were HLA class II+ inflammatory or CD8+ T cells associated with MXA expression. The potential mechanism underlying these associations are unclear. However, in neither disease were CD8+ T cell densities or proportions associated with fibrosis. Our sample was small and CD8+ T cells have been linked with progressive renal disease in RAR and LuN (*31, 32*). However, recent data from mouse models of LuN suggest that some of these populations have an exhausted or even protective phenotype (*61, 62*). Indeed, both PD-1 and ICOS expression were wide-spread on our infiltrating T cells. Our data indicate that, within our cohorts, macrophages are more associated with renal damage than conventional T cell populations.

There were both similarities and differences in the distribution of immune cell infiltrates in LuN and RAR. In both diseases, there was a strong enrichment in the periglomerular space. In contrast, the distribution of inflammation within the tubulointerstitium was different in LuN and RAR. In LuN, inflammation was evenly distributed within the interstitium without peritubular enrichment. In contrast, in RAR, there was enrichment at the peritubular border with some biopsies also manifesting tubulitis. These data suggest that peritubular and tubular inflammation characterize some RAR biopsies while LuN is characterized by interstitial inflammation. In LuN and RAR, we did not observe an enrichment of specific immune cells in any tubulointerstitial compartment. This suggests that there are no strong immunological barriers within the tubulointerstitium dictating the evolution of inflammation.

Spatial immunology is a new and evolving field in which the imaging and computational tools are rapidly improving (*63*). Currently, there are still technical limitations. While DAPI nuclear segmentation is still more reliable than whole cell segmentation, current techniques still under call irregularly shaped nuclei such as those of myeloid cells (*64*). We dilated nuclear segmentations to capture cytoplasmic staining for annotation.

While this is a reliable strategy for assessing lymphocytes, it often fails to capture peripheral staining on large cells such as tubule cells. In part, we circumvented this limitation by segmenting whole tubules. However, CD138 expression by tubules likely led to some plasma cells being included in the tubular mask. Finally, we used a hierarchical decision tree for cell class assignment which is a well-established approach used by others (*47, 55*). However, this strategy uses predefined cell classes and can miss novel cell populations. Therefore, our analysis is only an approximation. Nevertheless, within these limitations, we provide novel insights into *in situ* immune states and conceptual frameworks for further investigation.

Other approaches have biases and limitations. Single cell sorting from renal samples has provided a picture of the overall immune landscape that informed our staining panel (*22, 46*). However, different immune cell populations are likely extracted at different efficiencies from tissue. Furthermore, some cells, such as plasma cells, survive poorly during extraction and handling (*65*). These and other factors likely make scRNA-Seq from renal tissue variable and inefficient. In the one available study focused on *in situ* immune cells in LuN, only about 150 CD45+ cells were obtained per biopsy (*22*). In contrast, we identified over 7,000 immune cells per biopsy with spatial coordinates on each cell. This richness of information per biopsy enabled a dissection of renal inflammation heterogeneity not possible with current scRNA-Seq techniques. Finally, our experiments were done on archived FFPE biopsies. Future studies, with larger numbers of patients from longitudinal registries, can provide clinical context for different *in situ* immune states.

Our data reveal that in both LuN and RAR, and across almost all patients, the foundation of *in situ* inflammation is innate immunity. This is particularly surprising for LuN which is a manifestation of the canonical systemic autoimmune disease, SLE. Our studies characterize the most prevalent immune cell populations. Adaptive cells are present *in situ* and there are likely important functional relationships between these and resident innate cell populations. Furthermore, the relationships between systemic adaptive autoimmunity and *in situ* immunity, are largely unexplored in humans (*66*). Future mechanistic studies in SLE of concurrent blood and tissue biopsy samples, coupled with clinical trials of adaptive immune cell targeted therapies, will begin to unravel the complex inter-relationships between innate and adaptive cell programs and those between systemic and *in situ* autoimmunity.

## Methods

### Tissue acquisition

We obtained 54 archival blocks of kidney biopsies preserved as formalin-fixed, paraffin-embedded (FFPE) from the University of Chicago Human Tissue Resource Center. Within this, 25 blocks were from LuN patients, 23 were from RAR patients, and 6 were from the normal renal tissue at the margins of resected renal cell carcinomas.

### FFPE tissue processing and staining

Five-micrometer FFPE tissue sections were cut and mounted on 22mm×22mm glass coverslips. The tissue coverslips were deparaffinized as follows: The paraffin embedding was removed from the tissue sections via 20-min incubation at 60°C. Coverslips were transferred into Xylene and sequentially immersed in a fresh Xylene solution two times, 5 min each; 100% ethanol two times 5 min each; 95% ethanol 5 min; 70% ethanol 5 min; 50% ethanol 5min; 30% ethanol 5 min and distilled water 5 min. Tissue coverslips were then treated with 1× citrate buffer, pH6 (diluted from 100× stock, Abcam ab93678) for 20 min in a high pressure cooker. After antigen retrieval, tissues were then stained with Akoya’s staining kit for PhenoCycler (Akoya Biosciences, SKU7000008) following their protocol.

### Marker panel creation

We designed a 42-marker immunofluorescence panel after conducting a literature review of relevant immune cell populations and those identified on the landmark scRNA-Seq study of LN patient biopsies^8^. All antibodies were first validated with immunofluorescence staining on human tonsil and kidney sections. Validated antibodies were then conjugated with DNA-barcodes using the conjugation kit (Akoya Biosciences, SKU7000009) and revalidated using single-stain CODEX and multicycle CODEX runs. Provided below is a table of primary antibodies including source, dilution and CODEX cycle:

| Target | Clone | Manufacturer | Catalog No. | Dilution | Cycle |
| --- | --- | --- | --- | --- | --- |
| BDCA1 | OTI2F4 | Novus Biologicals | NBP2-70345 | 1:200 | 1 |
| BDCA2 | polyclonal | Novus Biologicals | AF1376 | 1:200 | 2 |
| CD10 | EPR22867 | abcam | ab256494 | 1:200 | 10 |
| CD103(ITGAE) | EPR22590-27 | abcam | AB254201-1001 | 1:200 | 9 |
| CD11c | EP1347Y | abcam | ab216655 | 1:200 | 1 |
| CD138 | EPR6454 | abcam | ab226108 | 1:200 | 4 |
| CD14 | SP192 | abcam | AB230903-1001 | 1:200 | 3 |
| CD16 | D1N9L | Cell signaling | 72204SF | 1:200 | 7 |
| CD163 | EDHu-1 | Novus Biologicals | NB110-40686 | 1:200 | 13 |
| CD20 | L26 | abcam | ab236434 | 1:200 | 8 |
| CD21 | EP3093 | abcam | ab271855 | 1:200 | 11 |
| CD27 | BLR083G | abcam | ab272072 | 1:200 | 10 |
| CD3 | SP7 | abcam | ab205228 | 1:200 | 12 |
| CD31 | EP3095 | abcam | ab226157 | 1:200 | 11 |
| CD4 | EPR6855 | abcam | ab181724 | 1:200 | 6 |
| CD43 | SPM503 | Novus Biologicals | NBP2-34775 | 1:200 | 12 |
| CD45 | EP322Y | abcam | ab214437 | 1:200 | 9 |
| CD45 | 2B11 + PD7/26 | Novus Biologicals | NBP2-34287 | 1:200 | 9 |
| CD56 | OTI1G4 | OriGene | CF506208 | 1:200 | 6 |
| CD68 | KP1 | eBioscience | #14-0688-82 | 1:200 | 13 |
| CD69 | EPR21814 | abcam | ab234512 | 1:200 | 8 |
| CD8 | C8/144B | Invitrogen | MA5-13473 | 1:200 | 7 |
| CD86 | E2G8P | Cell signaling |  | 1:100 | 5 |
| Claudin1 | ab238949 | abcam | EPR121871 | 1:200 | 10 |
| COLIII | polyclonal | Proteintech | 22734-1-AP | 1:200 | 4 |
| FOXP3 | PCH101 | eBioscience | 14-4776-82 | 1:200 | 4 |
| GZMA | EPR20161 | abcam | ab251499 | 1:200 | 5 |
| GZMB | D6E9W | Cell signaling | 79003SF | 1:200 | 9 |
| GZMK | polyclonal | abcam | EPR24601-164 | 1:200 | 8 |
| HLA-DP,-DR,-DQ | CR3/43 | abcam | ab7856 | 1:200 | 11 |
| ICOS | D1K2T | Cell signaling | 39740SF | 1:200 | 6 |
| IFN-g | EPR21704 | abcam | ab231301 | 1:200 | 7 |
| IL-10 | JES3-9D7 | Invitrogen | 14-7108-81 | 1:200 | 2 |
| IL-10 | polyclonal | Novus Biologicals | AF-217-NA | 1:200 | 2 |
| iNOS | SP126 | abcam | ab239990 | 1:100 | 3 |
| Ki67 | B56 | BD | BDB556003 | 1:200 | 7 |
| MERTK | Y323 | abcam | ab271851 | 1:200 | 13 |
| MTOC | TU30 | abcam | ab27074 | 1:200 | 12 |
| MUC-1 | 955 | Novus Biologicals | NBP2-44658 | 1:200 | 10 |
| MXA | polyclonal | R&D | AF7946 | 1:200 | 1 |
| PD1 | D4W2J | Cell signaling | 63815SF | 1:100 | 2 |
| ROR-γ | polyclonal | ThermoFisher Scientific | PA534164 | 1:200 | 3 |
| SLAMF7 | E5C4M | Cell signaling | 66110 | 1:200 | 1 |
| T-bet | D6N8B | Cell signaling | #27112SF | 1:100 | 5 |
| TCR-delta | H-41 | Santa Cruz | sc-100289 | 1:200 | 4 |

### Image acquisition and processing

Sections were deparaffinized and stained our panel of 42 antibodies each conjugated to a unique oligonucleotide. Images of full biopsy sections were acquired on an Andor Dragonfly 200 Spinning Disk Confocal Microscope (0.1507μm pixel size). The staining patterns of our 42-marker panel was acquired through iterative staining with Alexa Fluor 488, Atto550, Cy5/AF647, and AF750 fluorophores conjugated to complementary oligonucleotides and imaging using the PhenoCycler platform described previously^1^. Tissue autofluorescence images were also acquired at each imaging wavelength.

ASHLAR was used to stitch image tiles into a full-section composite and align the resulting channels. Ashlar performance was visually checked across all samples. Areas with insufficient alignment were rejected from downstream analysis. After aligning all image channels, the first blank cycle of imaging was used for background subtraction and normalization of all stained images. First, each channel of the blank cycle was subtracted from the corresponding fluorescence channel in all imaging cycles. Each imaging wavelength has a different dynamic range, so the subtracted images were also divided by the standard deviation of the background image to standardize dynamic range across imaging wavelengths. After standardization relative to imaging wavelength, images were min-max normalized to the 99th percentile. After this preprocessing, instance segmentation of cell nuclei was performed using Cellpose 2.0. Cell body segmentations were approximated by dilating the nucleus segmentations by 7 pixels (1.05 μm). Mean fluorescence intensity (MFI) was calculated from each marker in the panel using the cell body pixel mask as reference.

### Decision tree classifier and cell mapping to tissue

A subsample of 26 of the original 42 immunofluorescence markers were used to classify approximately 1.77 million cells. We used a decision-tree classifier for the multiclass annotation of cells that is analogous to flow-cytometry-based cell analyses and immunophenotyping. The decision tree considers the known experimental covariance between markers (i.e. CD3-expression would preclude CD4+ expression in T-lymphocytes). Cell positivity is determined by applying the multiotsu thresholding method using each cell’s MFI, across all 1.77 million cells in all 3 cohorts simultaneously. The multiotsu threshold that most closely matched manual spot validation was used. This manual spot validation was done by manually circling 5 positive signal cells in the desired cell population (ie, CD8+ expression for cytotoxic lymphocytes) and manually calculating the cell’s MFI in ImageJ. We further use our kidney control to check if cell thresholds are not under-calling or over-calling cells. Once classified, cells are mapped back to computational segmentations of the renal tissue for further downstream spatial analyses.

### Distinct immune trajectory

To statistically test for the differential presence of particular cell classes, we perform a non-parametric Mann-Whitney-U test for population differences using cell densities. We perform the following comparisons: LuN-KC, RAR-KC, and LuN-RAR. Benjamini-Hochberg p-value correction was performed to control for multiple p-value hypothesis testing.

### Renal structure segmentation

For the instance segmentation of kidney tubules, Omnipose was trained on 32 tiles of 10-x downsized DAPI kidney images (160×160 pixels) randomly selected from three kidney biopsies. The total number of training annotations was 355 instances. The change in training image size compared to the training image size in cellular segmentation was to reflect the size differences between a cell and a much bigger tubule structure. The training parameters were as follows: 1000 epochs, learning rate = 0.1, batch size = 16, number of classes = 2, tyx tuple input = 128×128. This model was trained from scratch without using any pretrained models. Post-training validation was done on 16 tiles of downsized DAPI kidney images (160×160 pixels), containing 195 tubule instances.

Whole-slide segmentation of lupus kidney biopsies was performed on DAPI channel after pre-processing and downsizing by a factor of 10. The segmentation parameters are as follows: mask threshold = 2.04, diameter = 30, with affinity segmentation. Post-segmentation of tubules was done to remove false positives (red blood cells clusters, clusters of aggregated lymphocytes in the interstitial spaces) by calculating the mean fluorescence intensity of canonical kidney structural markers per tubule object and subsequently removing the objects which express low tubule markers (CD10, MUC-1) and high expression level of all markers, which is typical of red blood cells. Using normalized mean fluorescence intensity, objects satisfying the following conditions are removed as false positives:

*μ*_CD10_ < 0.8, *μ*_MUC-1_ < 0.8, *μ*_Claudin-1_ < 1, *μ*_CD138_ < 1, 0.25 < (μMUC-1)/μCD10 <6.5

### DBSCAN

We used DBSCAN to find cellular clusters or neighborhoods in our multiplex microscopy imaging data. We used bootstrapping to subsample 75% of the DBSCAN data for a total of 3,000 repetitions for the ideal K number of clusters. Using the average bootstrap sum of squared distance plot and the delta sum of square distance plot we find that the empirically best fit K-clusters was around 9-11. We use 11 as optimal K for downstream analysis; moving forward with phenotyping only those cell neighborhoods with 20 and more cell members. To phenotype these DBSCAN segmented cell neighborhoods, we perform feature extraction by characterizing each neighborhood using the cell class count and total proportion for each of our 33 immune cell classes; we included the total cell count as another descriptor. Afterwards, to find unique defining features for each of our clusters, we generated a heatmap of the leave-one-out Z-test for every cluster.

### Computational resources

All computational tasks were carried out on the MEL server located in the Radiomics and Machine Learning Facility at the University of Chicago. MEL is equipped with 256 Xeon Gold 6130 CPU cores, 3 TB of DDR4 ECC RAM, 24 TB NVMe SSD storage space, and houses 16 Nvidia Tesla V100 32GB GPU accelerators.

---

**Supplemental Figure 1.**
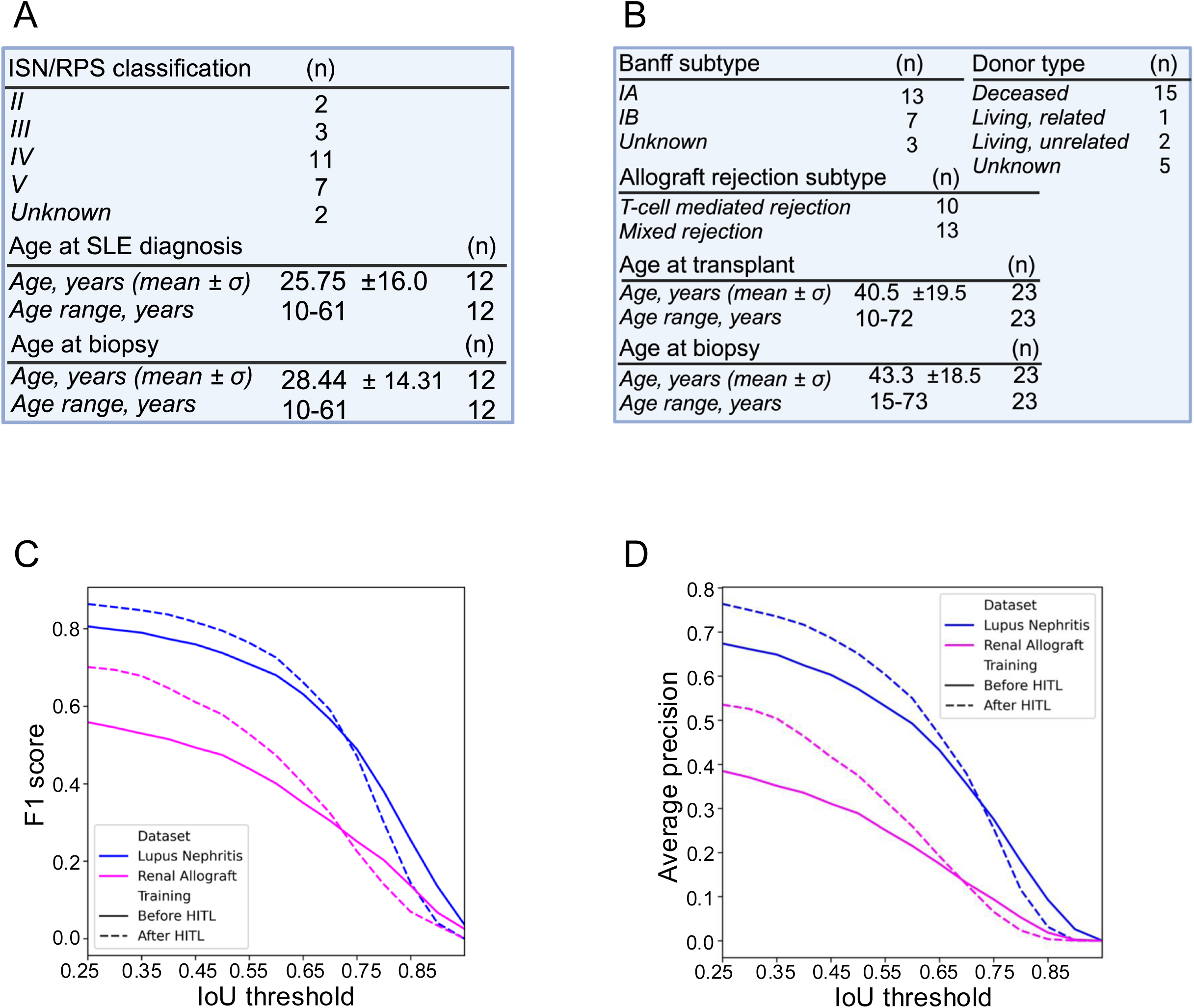
Cohort demographics and Cellpose HITL retraining. A. Summary table of patient descriptors and characteristics for lupus nephritis patients. Includes ISN/RPS nephritis class 2-5. B. Summary table of patient descriptors and characteristics for renal allograft rejection patients. Includes numbers with T-cell mediated rejection and mixed rejection. Other types of allograft rejection, such as antibody-mediated rejection, were excluded. C. Cellpose F1-score performance before Human-in-the-loop retraining (solid line) and after HITL retraining (dashed line). Lupus nephritis is shown in blue. Renal allograft rejection is shown in magenta. D. Cellpose Average Precision performance before Human-In-The-Loop retraining (solid line) and after HITL retraining (dashed line). Lupus nephritis is shown in blue. Renal Allograft Rejection is shown in magenta.

**Supplemental Figure 2.**
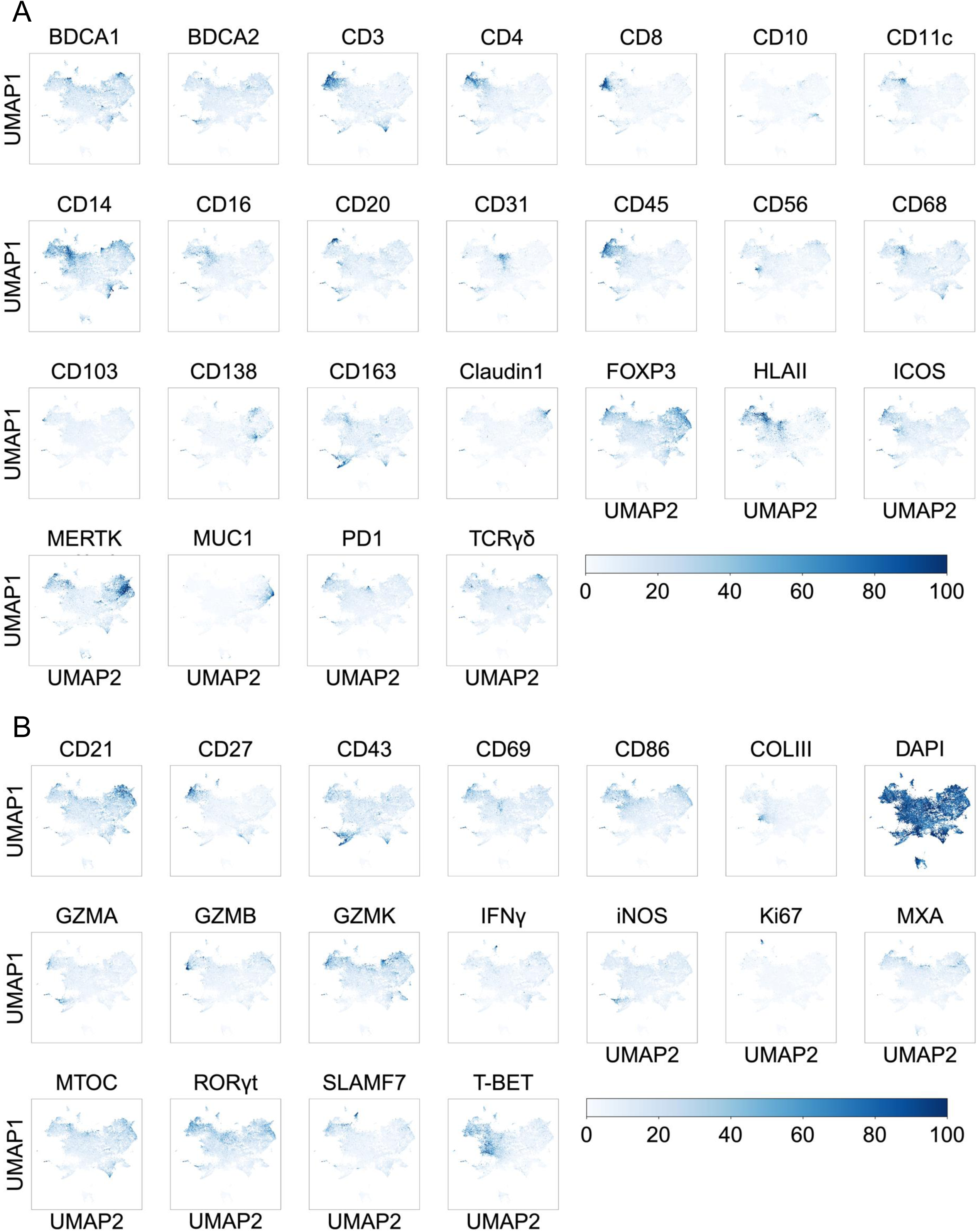
Average MFI expression of the 42-marker panel. A. UMAP dimensional reduction of cell body MFI from 30,000 cells randomly sampled. 10,000 cells are sampled from each of the three cohorts: KC, LuN, RAR. Shown are markers used for assigning cell class. MFI color scale is shown. B. UMAP dimensional reduction of cell body MFI as described prior. Shown are markers not used for cell class assignment.

**Supplemental Figure 3.**
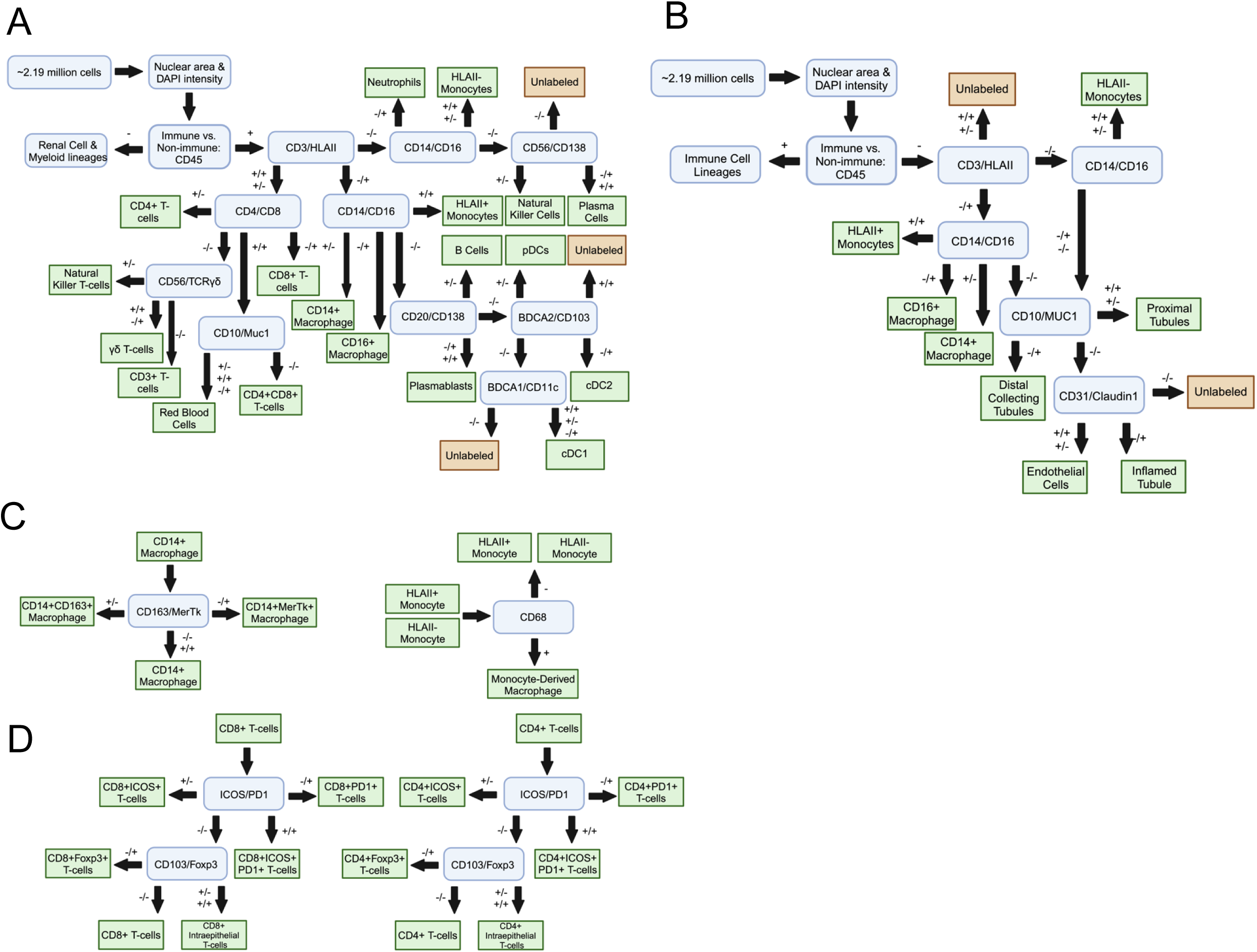
Decision trees for cell class assignment. A. Flow-cytometry analogous decision tree gating for immune cell classification based on cell body MFI. B. Flow-cytometry analogous decision tree gating for non-immune cell, and CD45 low/negative myeloid cell, classification based on cell body MFI. C. Decision tree gating for CD14+ macrophages (*left*) and monocytes (*right*). D. Decision tree gating for CD8+ T-cells (*left*) and decision tree gating for CD4+ T-cells (*bottom right*).

**Supplemental Figure 4.**
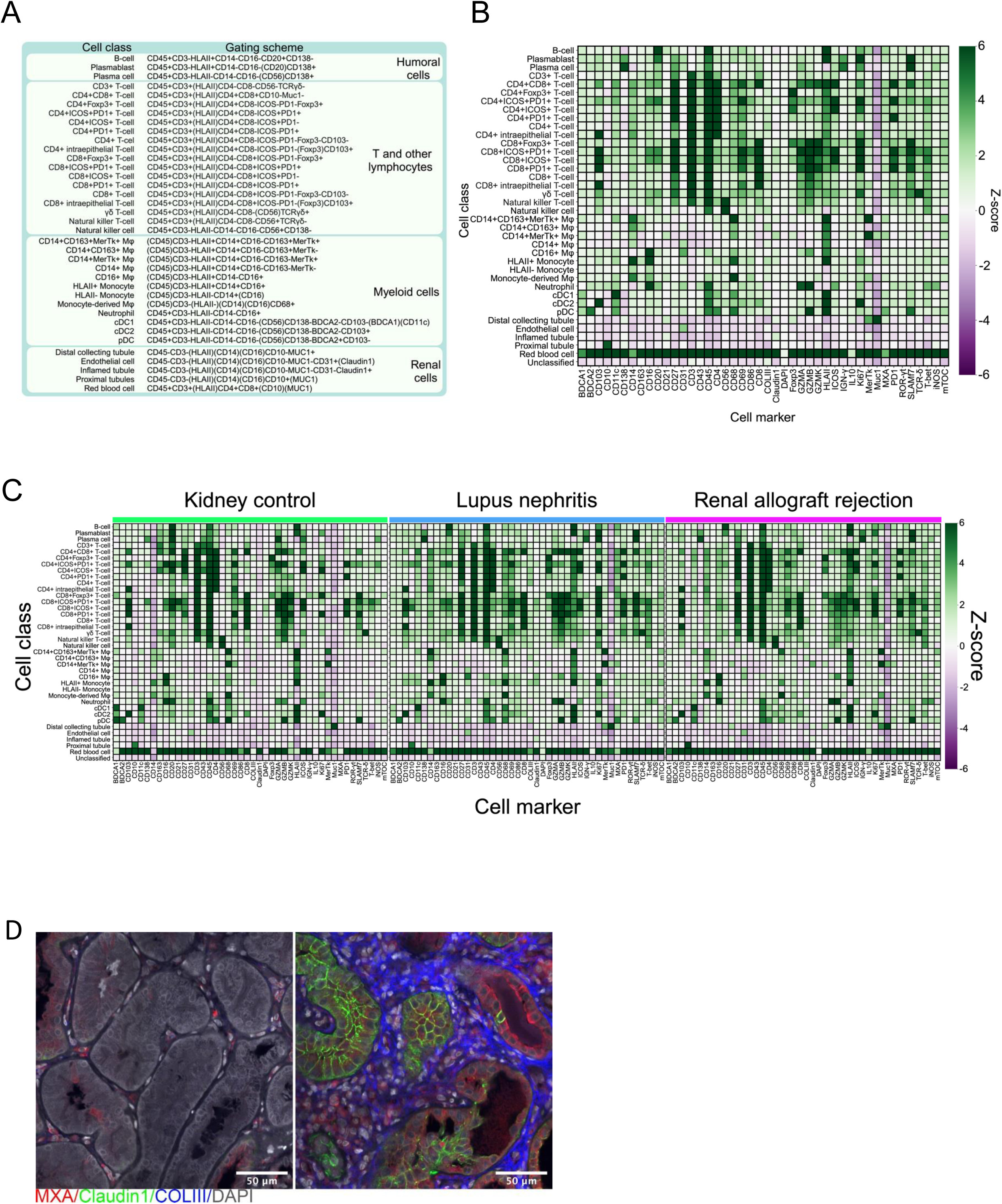
Cell classification of Cellpose segmented cells. A. Summary table of the 33 immune and 5 non-immune cell classes ultimately assigned using the decision tree algorithm; cells are grouped by lineage. B. Heatmap of the leave-one-out Z score (current cell class vs. all others) of the cell body MFI for the cell markers used in cell class assignment for the main immune and non-immune cell classes. C. Heatmap of the cohort leave-one-out Z score (current cell class vs. all others) of the cell body MFI for the cell markers used in cell class assignment. Heatmaps for kidney control (*left*), lupus nephritis (*middle*), and renal allograft rejection (*right*). D. Examples of normal, non-inflamed kidney tubules (left) and inflamed tubules expressing Claudin and MXA surrounded by COLIII fibrosis (right).

**Supplemental Figure 5.**
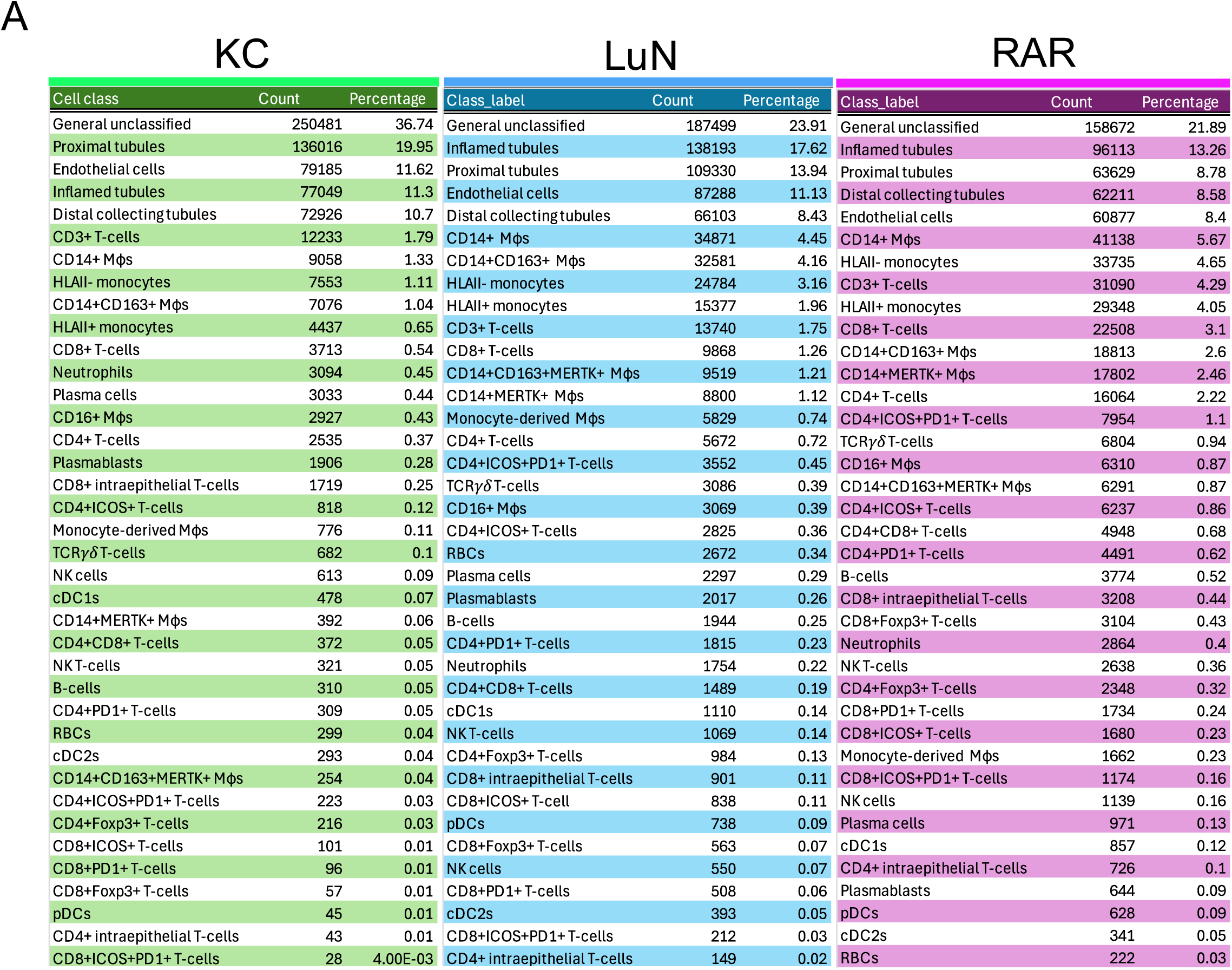
Cell class frequency data. A. Summary table of cell class total count and total percentage by disease cohort.

**Supplemental Figure 6.**
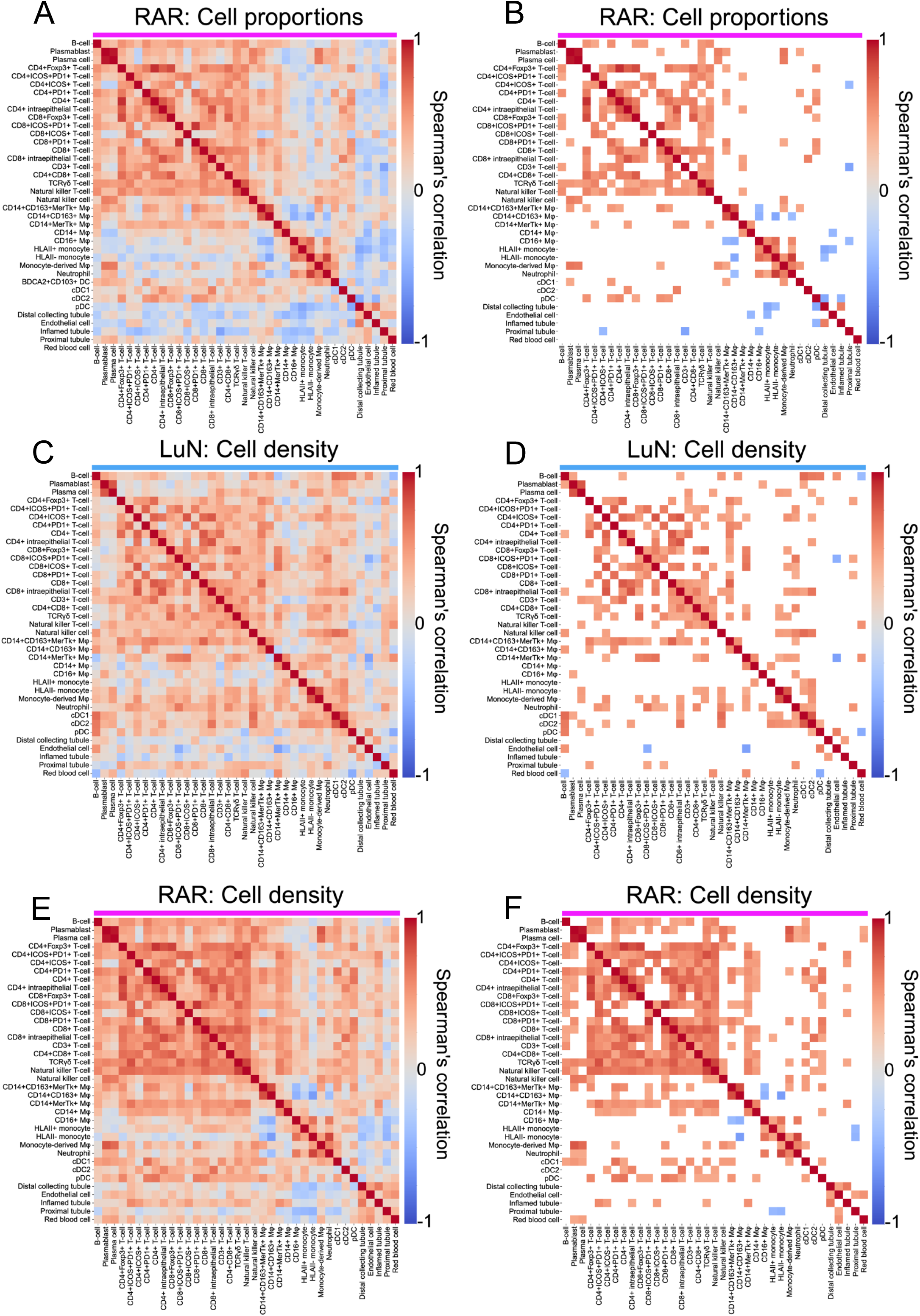
Cell Class Density and Proportion Correlations. A. Heatmap of the non-parametric spearman’s correlations between patient-level immune cell and non-immune cell proportions for renal allograft patients. B. Heatmap of the non-parametric spearman’s correlations between patient-level immune cell and non-immune cell proportions for renal allograft patients. Only significant correlations (p<0.05) are shown. C. Heatmap of the non-parametric spearman’s correlations between patient-level immune cell and non-immune cell density for lupus nephritis patients. D. Heatmap of the non-parametric spearman’s correlations between patient-level immune cell and non-immune cell density for lupus nephritis patients. Only significant correlations (p<0.05) are shown. E. Heatmap of the non-parametric spearman’s correlations between patient-level immune cell and non-immune cell density for renal allograft patients. F. Heatmap of the non-parametric spearman’s correlations between patient-level immune cell and non-immune cell density for renal allograft patients. Only significant correlations (p<0.05) are shown.

**Supplemental Figure 7.**
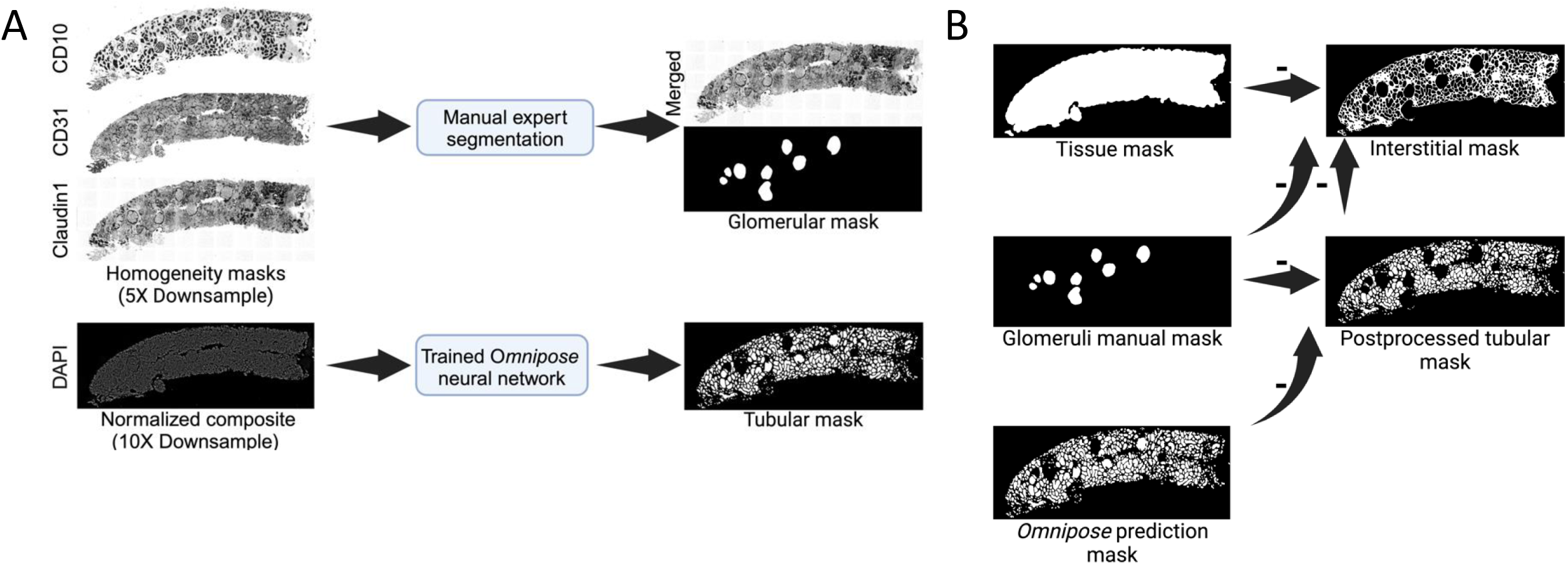
Kidney compartment segmentation workflow. A. Workflow of the procedure adopted for acquiring the computational segmentation of tubules and glomeruli. B. Procedure adopted for the creation of the interstitial mask and post-processed tubular mask.

**Supplemental Figure 8.**
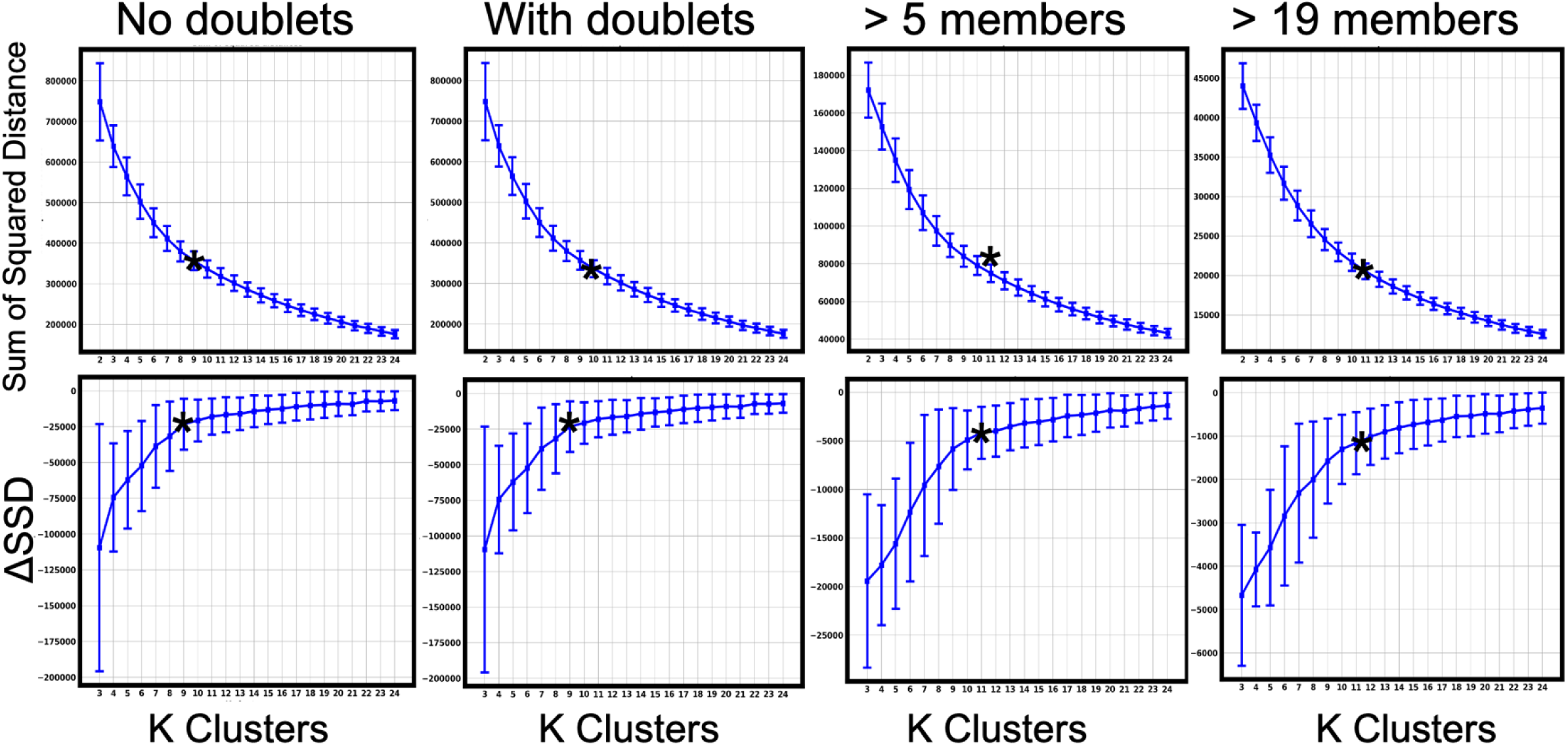
Estimation of the optimal cluster number. DBSCAN cell networks Bootstrap estimate of optimal K Clusters DBSCAN cell networks using: no doublets, with doublets, and above 5 cell members, and above 19 cell members respectively. Points are the average of 3000 repetitions with corresponding error bars. (Bottom) Δ sum of squared distances, and (top) sum of squared distances is shown. Asterisk indicates heuristic K optimal clusters for respective bootstrap experiment.

**Supplemental Figure 9.**
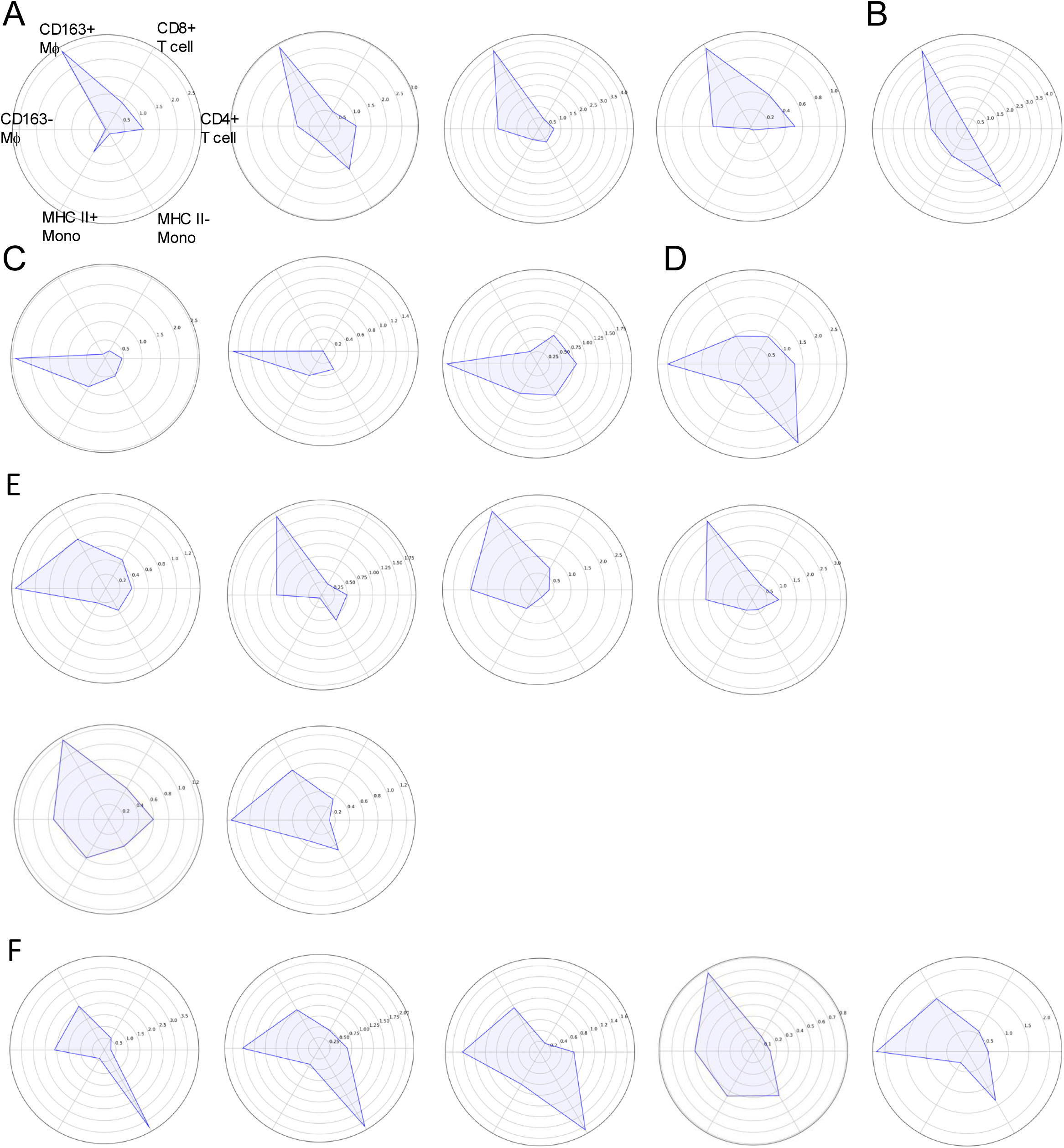
Principal immune cell trajectories of LuN biopsies. A. Examples of individual LuN biopsies with predominant CD163+ macrophage trajectories. B. Example of LuN biopsy with predominant CD163+ macrophage and HLA II-inflammatory monocyte trajectories. C. Examples of LuN biopsies with predominant CD163-macrophage trajectories. D. Example of LuN biopsy with predominant CD163-macrophage and HLA II-inflammatory monocyte trajectories. E. Examples of LuN biopsies with both CD163+ and CD163-macrophage trajectories. F. Examples of LuN biopsies with CD163+ and CD163-macrophage trajectories as well as HLA II-inflammatory monocyte trajectories.

**Supplemental Figure 10.**
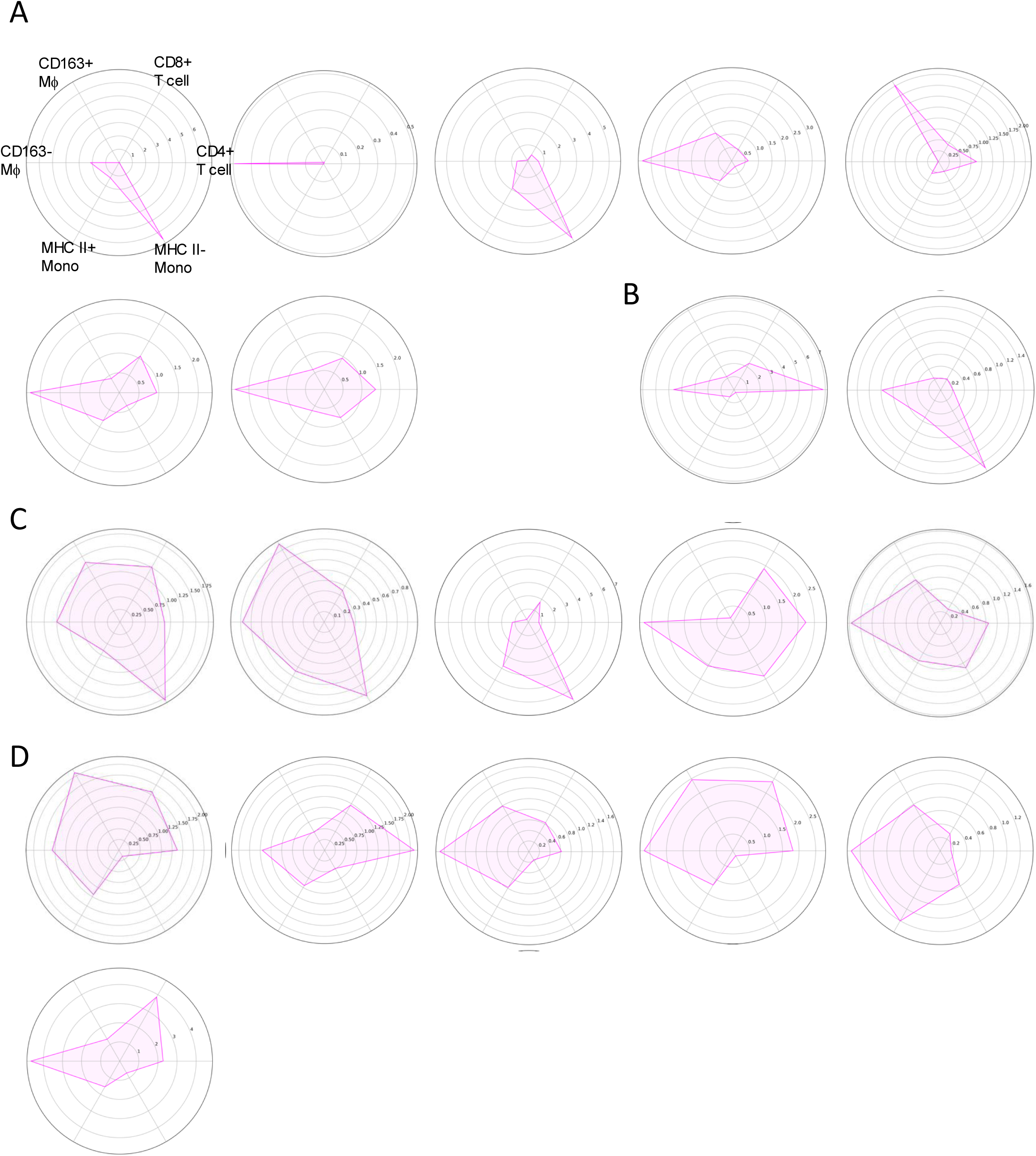
Principal immune cell trajectories of RAR biopsies. A. Examples of RAR biopsies with predominance of a single immune cell trajectory. B. Examples of RAR biopsies with two immune cell trajectories. C. Examples of RAR biopsies with multiple immune cell trajectories including HLA II-inflammatory monocytes. D. Examples of RAR biopsies with multiple immune cell trajectories excluding HLA II-inflammatory monocytes.

